# A probabilistic framework for cellular lineage reconstruction using single-cell 5-hydroxymethylcytosine sequencing

**DOI:** 10.1101/739300

**Authors:** Chatarin Wangsanuwat, Javier F. Aldeguer, Nicolas C. Rivron, Siddharth S. Dey

**Author notes:** Correspondence (S.S.D).

## Abstract

Lineage reconstruction is central to understanding tissue development and maintenance. While powerful tools to infer cellular relationships have been developed, these methods typically have a clonal resolution that prevent the reconstruction of lineage trees at an individual cell division resolution. Moreover, these methods require a transgene, which poses a significant barrier in the study of human tissues. To overcome these limitations, we report scPECLR, a probabilistic algorithm to endogenously infer lineage trees at a single cell-division resolution using 5-hydroxymethylcytosine. When applied to 8-cell preimplantation mouse embryos, scPECLR predicts the full lineage tree with greater than 95% accuracy. Further, scPECLR can accurately extract lineage information for a majority of cells when reconstructing larger trees. Finally, we show that scPECLR can also be used to map chromosome strand segregation patterns during cell division, thereby providing a strategy to test the “immortal strand” hypothesis in stem cell biology. Thus, scPECLR provides a generalized method to endogenously reconstruct lineage trees at an individual cell-division resolution.

## Introduction

Understanding lineage relationships between cells in a tissue is one of the central questions in biology. Reconstructing lineage trees is not only fundamental to understanding tissue development, homeostasis and repair but also important to gain insights into the dynamics of tumor evolution and other diseases. Genetically encoded fluorescent reporters have been a powerful approach to reconstruct the lineage of many tissues^1^. However, these methods require the generation of complex animal models for each stem or progenitor cell type of interest, and are limited to a clonal resolution^1^. Similarly, other pioneering techniques, such as the use of viruses^2^, transposons^3,4^, Cre-loxP based recombination^5^ and CRISPR-Cas9^6-8^ have also been used to genetically label cells to primarily reconstruct clonal lineages that lack the resolution of an individual cell division. This clonal resolution limits our ability to understand tissue dynamics at a single cell-division resolution. While a recent report that combined CRISPR-Cas9-mediated targeted mutagenesis with single-molecule RNA fluorescence *in situ* hybridization enabled reconstruction of lineages at a single cell-division resolution (MEMOIR)^9^, its ability to infer lineages dropped substantially by the 3^rd^ cell division.

Further, as all these methods involve exogenous labeling strategies, they cannot be used to map cellular lineages in human tissues directly, thereby posing a significant barrier to understanding human development and diseases. While endogenous somatic mutations have been used to reconstruct lineages, the low frequency of their occurrence and distribution over the whole genome make them challenging to detect and therefore limit their application as a lineage reconstruction tool^10-12^. Similarly, a recent method used mutations within the mitochondrial genome to reconstruct lineages, but as most other lineage reconstruction approaches, it is limited to a clonal resolution^13^. Previously, we developed a method to detect the endogenous epigenetic mark 5-hydroxymethylcytosine (5hmC) in single cells (scAba-Seq) and showed that the lack of maintenance of this mark during replication coupled with the low rates of Tet-mediated hydroxymethylation resulted in older DNA strands containing higher levels of 5hmC^14^. The ability to track individual DNA strands through cell division allowed us to deterministically reconstruct lineages that were limited to 2 cell divisions^14^. Therefore, to overcome limitations of existing methods, we report scPECLR (single-cell Probabilistic Endogenous Cellular Lineage Reconstruction), a generalized probabilistic framework for endogenously reconstructing cellular lineages at an individual cell division resolution using single-cell 5hmC sequencing.

## Results

### Genome-wide strand-specific 5hmC enables initial lineage bifurcation of individual cells into two subtrees

As proof-of-principle, we dissociated 8-cell mouse embryos and performed scAba-Seq to quantify strand-specific genome-wide patterns of 5hmC in single cells (Figure 1-I). As shown previously, a majority of 5hmC is present on the paternal genome during these stages of preimplantation development^15-17^. Single cells from an 8-cell embryo displayed a mosaic genome-wide distribution with no overlap of 5hmC between the plus and minus strands of a chromosome (Figure 1-II). Further, we found that for each chromosome, the strand-specific 5hmC was localized to a few cells with other cells containing undetectable levels of the mark (Figure 1-II). These observations clearly demonstrate that only one allele carries a majority of 5hmC, and that consistent with previous results, we are primarily detecting 5hmC on the original paternal genome with DNA strands synthesized in subsequent rounds of replication carrying very low levels of the mark. We used this as our basis to reconstruct cellular lineages of 8-cell mouse embryos.

**Figure 1.**
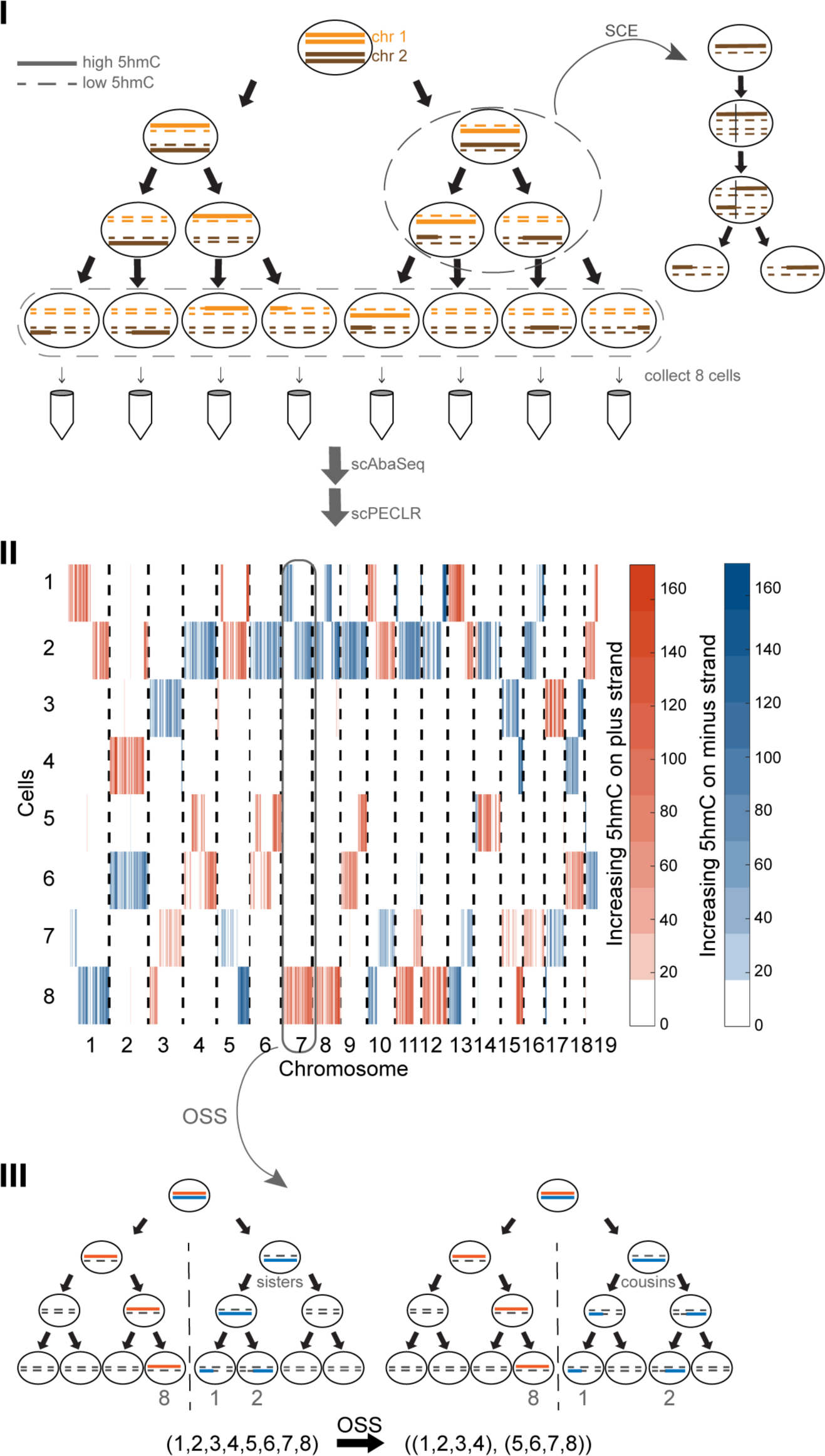
Strand-specific single-cell 5hmC data enables initial lineage bifurcation of individual cells into two subtrees. (I) Schematic shows a zygote with chromosomes containing high 5hmC levels (solid lines) undergoing three cell divisions. The newly synthesized strands at each cell division contain very low levels of 5hmC (dotted lines). SCE events occur randomly during each cell cycle. All cells are isolated and sequenced using scAba-Seq to quantify strand-specific 5hmC in single cells. (II) Data shows mosaic pattern of strand-specific 5hmC in single cells obtained from an 8-cell mouse embryo. 5hmC counts within 2 Mb bins on the plus and minus strands of all chromosomes are shown in orange and blue, respectively. (III) The original plus and minus strands of each chromosome should be found in cells on opposite sides of the lineage tree. This OSS analysis on chromosome 7 places cell 8 in one 4-cell subtree and cells 1 and 2 in the other subtree. Performing OSS on all chromosomes places cells in one of these two 4-cell subtrees and reduces the complexity of the lineage reconstruction problem.

As the first step towards reconstructing lineage trees, we noted that the original plus and minus strands of each paternal chromosome in the 1-cell zygote will be found in distinct cells on opposite sides of the lineage tree after *n* cell divisions. As a result, all cells can be placed in one of two subtrees, thereby reducing the number of cell divisions to be reconstructed from *n* to *n* − 1. For example, at the 8-cell stage, the original plus strand of chromosome 7 is detected in cell 8 while the minus strand is detected in cells 1 and 2 (Figure 1-II). This suggests that cell 8 is on the opposite side of the lineage tree compared to cells 1 and 2. Performing this first step of scPECLR, which we refer to as original strand segregation (OSS) analysis, over all the chromosomes enables us to systematically place cells 1-4 and 5-8 on opposite sides of the lineage tree for this embryo, reducing the complexity of the problem from reconstructing 3 cell divisions with 315 tree topologies to 2 cell divisions with 9 tree topologies (Figure 1-III).

### Probabilistic lineage reconstruction using scPECLR accurately predicts 8-cell embryo trees

To reconstruct the complete lineage tree, we next focused attention on the mosaic pattern of 5hmC arising from abrupt transitions in hydroxymethylation levels among cells along the length of a chromosome. As described previously, these sharp transitions in 5hmC that are shared between two cells are the result of homologous recombination during sister chromatid exchange (SCE) events in the G2 phase of a previous cell cycle^14^. Detection of 5hmC transition points that are common to two cells therefore indicate a shared evolutionary history between these cells (Figure 1-I, inset). However, while a SCE event at the 4-cell stage would imply that the cells are sister cells (Figure 1-III, left), one occurring at the 2-cell stage would indicate that the same pattern of 5hmC transition can also be observed between cousin cells (Figure 1-III, right). Thus, the observation of a single shared SCE event between two cells cannot be used to immediately discriminate between sister and cousin cell configurations.

To systematically determine the likelihood of observing different tree topologies, we developed a probabilistic framework where the occurrence of SCE events are modeled as a Poisson process. The total number of SCE events is used to estimate the parameter *b* of the Poisson process, the rate of SCE events per chromosome per cell division, using maximum likelihood estimation (Methods). Following OSS, 8-cell trees can be grouped into two 4-cell subtrees, each with 3 possible tree arrangements (Figure 2-I). Next, we used the probabilistic model to calculate the likelihood of observing a SCE pattern for a chromosome given a tree topology. We observed a large variety of SCE patterns, ranging from commonly observed patterns, such as one or two SCE transitions shared between two cells, to more complex distributions of 5hmC between cells (Supplementary Figure 1). For the most common pattern of one SCE transition between two cells, scPECLR predicts that the tree with the two cells as sisters (Tree A) is twice as likely as one where the two cells are cousins (Tree B or C), in good agreement with simulated data (Figure 2-II and Methods). Similarly, both our model prediction and simulations show that when two SCE transitions are shared between two cells, the probability that the two cells are sisters is 2 to 3 times higher than the probability that they are cousins, with the likelihood ratio between sister and cousin tree configurations depending on the relative position of the SCE transition on the chromosome (Figure 2-III and Methods). The lower probability of observing this pattern in cousin tree arrangements arises from the constraint that only even number of SCE transitions can occur within the region between *k*_11_ and *k*_13_ of the chromosome during the last cell division (Supplementary Figure 2). More complex 5hmC distribution patterns, such as when two SCE events are shared between three cells substantially favors the Tree A configuration (Figure 2-III and Methods). After the SCE pattern of each chromosome is analyzed, we can estimate the total likelihood of observing the different tree topologies, assuming that the SCE events on each chromosome are independent (Methods). Finally, the likelihood of an 8-cell tree is the product of the likelihoods of the two corresponding 4-cell subtrees (Figure 2-IV).

**Figure 2.**
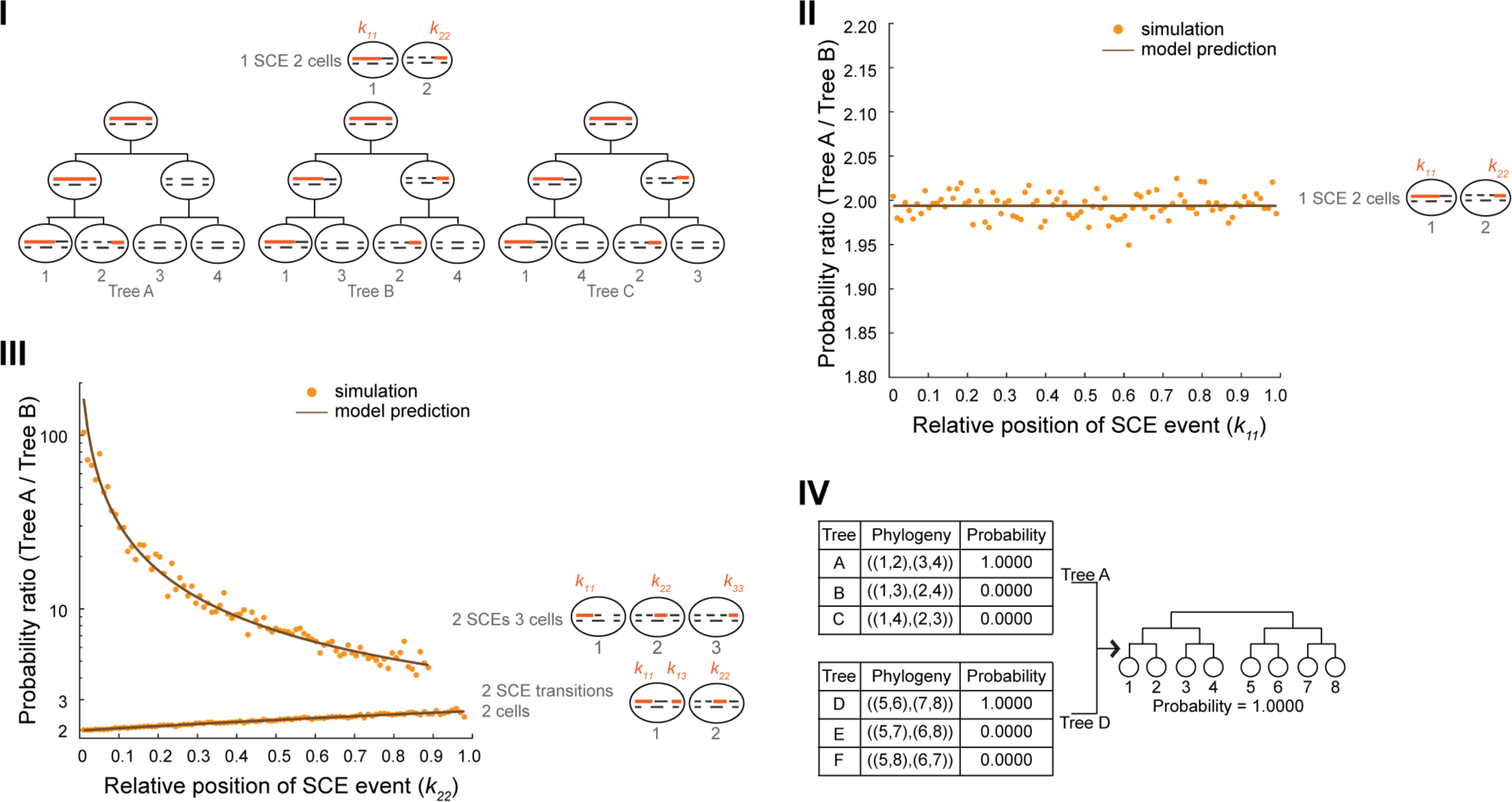
Endogenous 5hmC based lineage reconstruction using scPECLR. (I) Schematic showing that two cells sharing an original DNA strand (solid orange line) can either be sisters (Tree A) or cousins (Trees B and C) depending on whether the SCE event occurred at the 4- to 8- or 2- to 4-cell stage, respectively. All newly synthesized DNA strands are shown as dashed black lines. (II) For the case of a SCE transition between two cells, the probability of the pair of cells being sisters *vs*. cousins is plotted against the relative position of the SCE event on the chromosome (*k*_11_). The model prediction (black) and simulation results (yellow) are shown for chromosome 1 (*N* = 97 for 2 Mb bins) with *b* = 0.3. (III) The probability ratio between Trees A and B are shown as a function of *k*_22_ for *N* = 97 and *b* = 0.3 for two cases – 2 SCE transitions shared between 2 cells and 2 SCE events shared between 3 cells. (IV) For the 8-cell mouse embryo shown in Figure 1-II, the probability of observing the different topologies of the two 4-cell subtrees are shown.

To test the accuracy of scPECLR, we simulated 5hmC patterns of 8-cell embryos with a SCE rate similar to the experimentally observed value (*b* = 0.3), which is also within the range of SCE event rates found in various other cell types^18-22^. We found that scPECLR predicted the lineage tree correctly in 96% of all simulations (Figure 3-I). In contrast, MEMOIR predicted the lineage tree accurately in only ∼67% of the top 40% most reliably reconstructed trees (Figure 3-I). This improved accuracy strongly suggests that endogenous strand-specific 5hmC patterns present an accurate tool to reconstruct lineage trees at an individual cell division resolution. An automated pipeline to reconstruct cellular lineages is provided with this work (Methods). We next applied scPECLR on the 8-cell mouse embryo shown in Figure 1-II and other embryos to predict lineage trees with high confidence (Figure 2-IV and Supplementary Figure 3).

**Figure 3.**
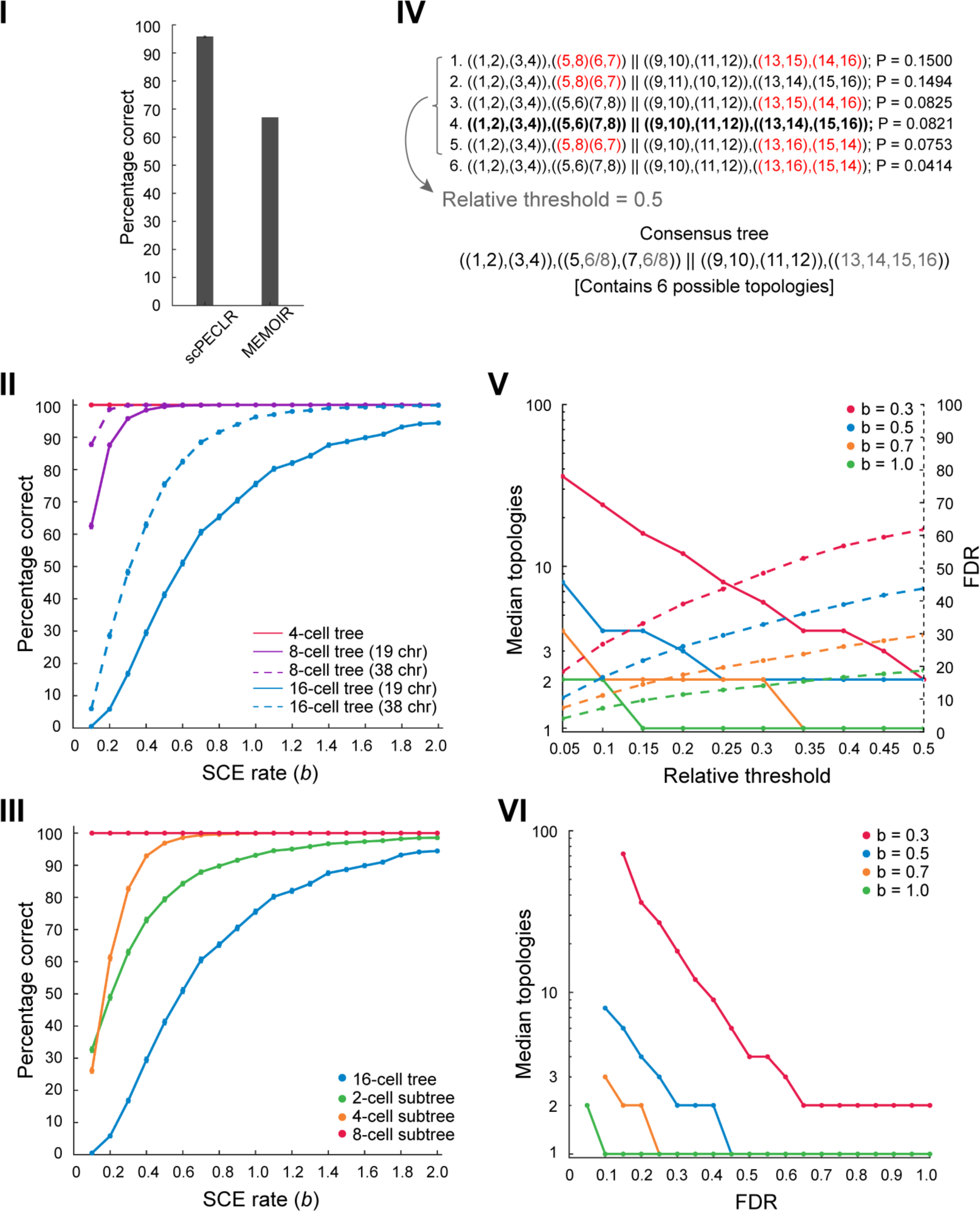
scPECLR can reconstruct 8-cell lineage trees accurately and be extended to reconstruct larger lineage trees. (I) scPECLR accurately predicts the lineage of 96% of simulated 8-cell trees (*b* = 0.3). Error bars indicate the bootstrapped standard error. In comparison, MEMOIR accurately predicts 67% of the top 40% most reliably reconstructed 8-cell trees^9^. (II) Panel shows the percentage of simulated 8-cell and 16-cell trees that are correctly predicted by scPECLR for different SCE rates (*b*). Solid and dotted lines indicate cells where 5hmC can be quantified in 19 or 38 chromosomes, respectively. The prediction accuracy is computed by simulating 5000 trees. Error bars indicate the bootstrapped standard error of prediction accuracy. (III) Panel shows the percentage of 2-, 4- and 8-cell subtrees that are accurately predicted within simulated 16-cell trees as a function of the SCE rate (*b*). The prediction accuracy is computed by simulating 5000 16-cell trees. Error bars indicate the bootstrapped standard error of prediction accuracy. (IV) Schematic illustrating how consensus trees are obtained. In this example, the top 6 tree topologies (with the highest probabilities) that are obtained after applying scPECLR on a simulated 16-cell tree are shown. The relative threshold parameter is used to determine the number of topologies that are considered in the consensus tree analysis. With a relative threshold of 0.5, the top 5 tree topologies in this example are used to generate a consensus tree that is consistent with all these trees. The uncertainty within the consensus tree is quantified by the number of tree topologies it contains. The higher the number of tree topologies it contains, the higher is the uncertainty within the consensus tree. Red fonts indicate parts of the lineage tree that are incorrectly predicted. The tree highlighted in bold is the true tree. (V) Simulation results show that as the relative threshold increases, the median number of topologies in the consensus tree decreases (solid lines, left axis), while the false discovery rate (FDR) increases (dotted lines, right axis). For these simulations, two other parameters *t*8 and *t*4 are set to 0.75 and 1.0, respectively. (For additional details on the parameters, see Methods). (VI) As the FDR decreases, the median number of topologies contained within the consensus tree increases. Thus, this panel shows how the specificity of the consensus tree is related to error tolerance. For *b* ≥ 0.7, the median number of topologies contained within the consensus tree rapidly drops to 1, suggesting that the consensus tree is fully constrained and is the correct tree. Note, the lowest FDR possible for *b* = 0.3, 0.5, 0.7 and 1.0 are 15%, 10%, 10%, and 5%, respectively.

As SCE transitions play a central role in reconstructing cellular lineage trees with scPECLR, we next explored how the endogenous rate of SCE events influences the accuracy of the model. As expected, the accuracy of lineage reconstruction increases monotonically with increasing rates of SCE events, with greater than 98% of the simulated 8-cell trees correctly predicted for *b* ≥ 0.4 (Figure 3-II and Methods). These simulations were performed using 19 paternal autosomes, consistent with our observation that a majority of 5hmC is found on the paternal genome in preimplantation mouse embryos. However, most cell types carry 5hmC on both parental genomes and therefore, we also performed simulations with 38 chromosomes. Again, as expected, the predictive power of the model increases, with the lineages of more than 98% of the simulated 8-cell trees accurately predicted for *b* ≥ 0.2 (Figure 3-II). These results demonstrate that the lineage tree can be accurately predicted up to 3 cell divisions even with low rates of SCE events (Figure 3-II).

### scPECLR can be extended to reconstruct larger lineage trees

We next extended this approach to reconstruct the lineage of 16-cell trees, where the number of possible tree topologies increase exponentially to more than 6×10^8^. While the ability to predict the complete lineage tree decreases (17% of all simulated 16-cell trees were predicted correctly for *b* = 0.3), we found that in a majority of cases large parts of the lineage tree were reconstructed accurately with the most common error being the misidentification of one sister pair within a 4-cell subtree (Figures 3-II and 3-III). For a SCE rate of *b* = 0.3, 83% of all 4-cell subtrees and 63% of all 2-cell subtrees (sister pairs) were predicted correctly (Figure 3-III). Overall, when reconstructing a 16-cell lineage tree, over 80% of all 4-cell and 2-cell subtrees were predicted accurately for *b* ≥ 0.3 and *b* ≥ 0.6, respectively (Figure 3-III). These results suggest that when reconstructing 16-cell trees from strand-specific 5hmC data, it will be important to identify parts of the lineage tree that we can predict with high confidence. To accomplish this, we first included all tree topologies that were predicted to have probabilities above a threshold relative to the tree with the highest probability (Figure 3-IV). A consensus tree that is consistent with all these tree topologies is then established (Figure 3-IV, Supplementary Figure 4 and Methods). For example, with *b* = 0.3 and for a relative threshold of 0.1, the median consensus tree contained 24 tree topologies (Figure 3-V, solid red line). The consensus trees displayed a false discovery rate (FDR) of ∼26%, implying that in 26% of the simulations, the consensus tree was not consistent with the true tree (Figure 3-V, dotted red line). As the relative threshold is increased (that is, we include fewer tree topologies to construct the consensus tree), the median consensus tree contains fewer topologies, resulting in a more specific or constrained consensus tree. However, this comes at the expense of an increase in FDR. Thus, the relative threshold allows us to tune the competing goals of specificity and accuracy of the consensus tree. These results show that for a certain rate of SCE events and a desired level of FDR, the median number of topologies contained in the consensus tree can be estimated, yielding insights into how much lineage information can be extracted from the 5hmC data based on the number of SCE events and our error tolerance (Figure 3-VI and Methods).

### scPECLR can be used to infer the rate of SCE events at each cell division and test the “immortal strand” hypothesis

In addition to reconstructing cellular lineage trees, scPECLR can also be used to infer the rate of SCE events at individual cell divisions. For example, for 8-cell mouse embryos, the 5hmC distribution at the 4-cell and 2-cell stages can be reconstituted based on the predicted lineage, enabling us to estimate the rate of SCE events at each cell division (Figure 2-IV and Supplementary Figure 3). While the overall SCE rate over three cell divisions for all the 8-cell mouse embryos analyzed in this study was estimated to be 0.31 events per chromosome per cell division, the individual SCE rates for the 1-to-2, 2-to-4, and 4-to-8 cell stages were 0.13, 0.11, and 0.68, respectively. Further, we found that the different rates of SCE events at each cell division did not affect the prediction accuracy of scPECLR (Supplementary Figure 5 and Methods). These results show that scPECLR can be used to infer the rate of double-stranded DNA breaks at each cell division and that the rate of SCE events can vary during development.

Finally, we explored another application of scPECLR. As scPECLR uses endogenous strand-specific 5hmC in single cells to reconstruct 8-cell trees with high accuracy, we hypothesized that this method could be used to quantify how parental alleles are segregated during cell division (Figure 4-I). Different stem cell populations, such as hair follicle^23^, neural^24^, satellite muscle^25,26^ and intestinal crypt stem cells^27,28^, have previously been shown to display non-random segregation of DNA strands that can influence cell fate decisions. These results have led to the “immortal strand” hypothesis that postulates old DNA strands are retained by daughter stem cells during asymmetric cell divisions to reduce the mutational load arising from genome replication of these longer lived cells. During mouse preimplantation development, recent reports have shown that blastomeres show biases in cell fate specification as early as the 4-cell stage^29,30^. Therefore, as proof-of-concept, we investigated sister chromatid segregation patterns at the 4-cell stage. To do this, we first combined 5hmC data from reconstructed sister cell pairs at the 8-cell stage to generate the distribution of the oldest DNA strands at the 4-cell stage (Figure 4-II). In the example shown, when comparing cells (1,2) and (3,4), we found that the original DNA strands preferentially segregate to cell (1,2). In contrast, such a non-random pattern of DNA strand segregation is not observed between sister cells (5,6) and (7,8). Quantitatively, we analyzed seven mouse embryos to find one sister pair at the 4-cell stage that displayed statistically significant non-random segregation of DNA strands (Figure 4-III and Methods). Thus, this proof-of-concept study shows that strand-specific reconstruction of lineage trees can be a powerful approach to test the immortal strand hypothesis in different stem cell populations.

**Figure 4.**
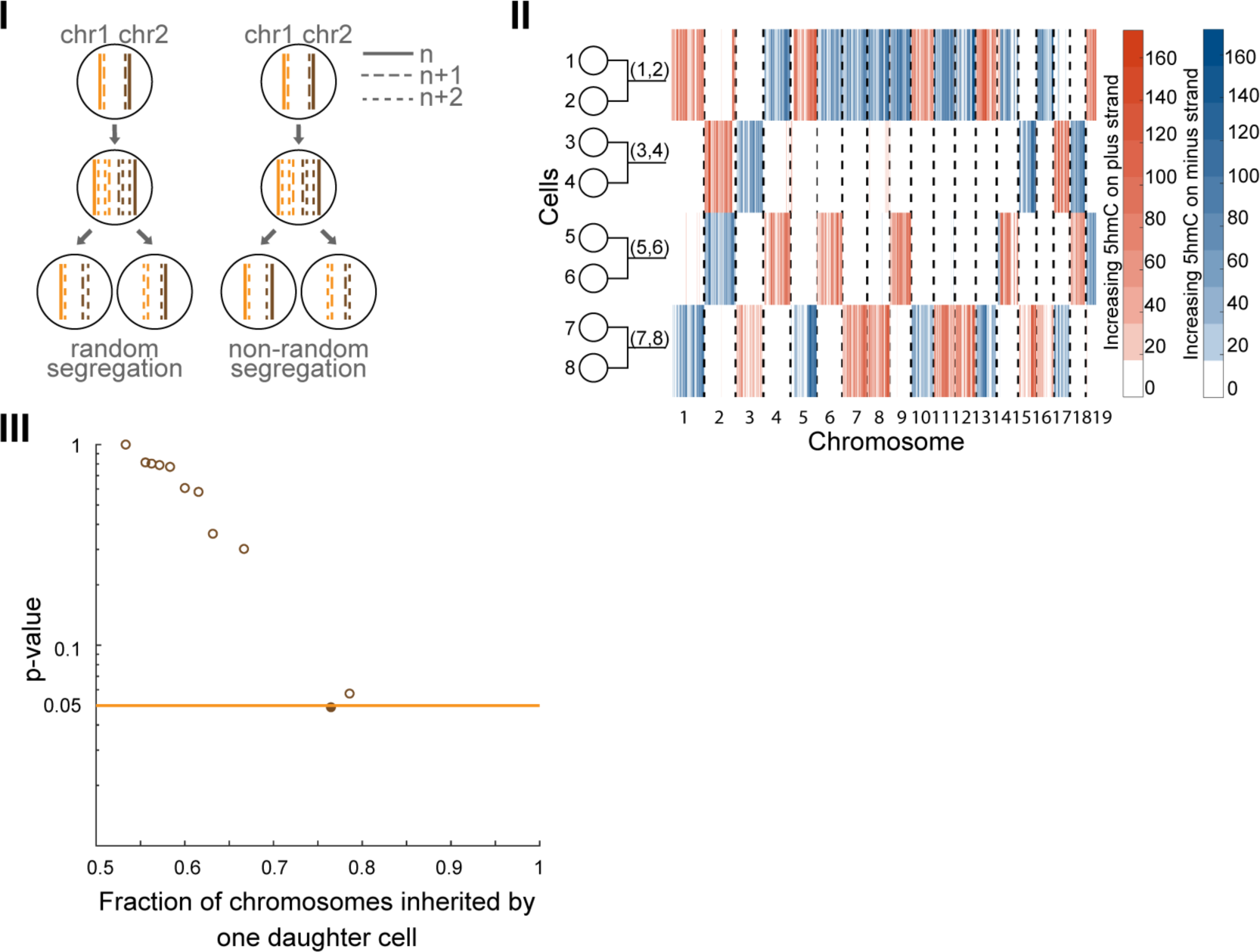
scPECLR can be used to map DNA strand segregation patterns. (I) Schematic showing DNA strand segregation patterns during cell division. Non-random segregation results in old DNA strands being preferentially inherited by one daughter cell. The oldest DNA strands are shown as solid lines, and strands synthesized in the *n* + 1 and *n* + 2 generation are shown as dashed and dotted lines, respectively. (II) Combining the experimental 5hmC data for the 8-cell embryo in Figure 1-II with the lineage tree predicted by scPECLR enables the genome-wide reconstitution of 5hmC in single cells at the 4-cell stage. (III) Proof-of-principle testing non-random segregation of DNA strands at the 4-cell stage of mouse embryogenesis. The *p*-values from a binomial test under a null hypothesis of random segregation shows that one embryo displays statistically significant (*p* < 0.05) non-random segregation of DNA strands.

## Discussion

Cellular lineage reconstruction plays an important role in answering fundamental questions in several areas of biology, such as immunology, cancer biology and developmental and stem cell biology. However, most current methods have two major limitations: (1) Clonal lineage reconstruction that cannot establish lineage relationships at the resolution of individual cell divisions; and (2) The use of transgenes that involves the time-intensive generation of complex animal models and is an approach that cannot be extended to map lineages in human tissues. To overcome these limitations, we have developed a generalized probabilistic framework to reconstruct cellular lineages at an individual cell division resolution using strand-specific single-cell 5hmC sequencing data. scPECLR can potentially also be combined with single-cell measurements of other non-maintained epigenetic marks, such as 5-formylcytosine (5fC) and 5-carboxylcytosine (5caC), to reconstruct lineages^21^. Importantly, the use of an endogenous epigenetic mark to reconstruct lineage trees suggests that this method can be directly extended to study human development.

In future, combining detection of 5hmC with measurements of mRNA from the same cell can potentially be used to simultaneously quantify both the cell type and the lineage relationship between cells in a tissue, thereby enabling us to directly probe symmetric and/or asymmetric cell fate decisions of stem cells at an individual cell division resolution. Such measurements can provide detailed insights into how stem cells maintain an exquisite balance between self-renewal and differentiation to regulate the dynamics of tissue development and homeostasis. Finally, we anticipate that integrating 5hmC based lineage reconstruction with measurements of other epigenetic marks from the same cells holds tremendous promise in understanding the genome-wide transmission and inheritance of the epigenome at each cell division.

## Supporting information

Code (scPECLR)

## Acknowledgement

We thank Alex Chialastri and other members of the Dey lab for helpful feedback. We acknowledge support for the computational work from the Center for Scientific Computing at the California NanoSystems Institute (CNSI) and Materials Research Laboratory (MRL) at UCSB: an NSF MRSEC (DMR-1720256) and NSF CNS-1725797. This work was supported by a CNSI Challenge Grant (8-447810-69085).

## Author contributions

C.W. and S.S.D. designed the study. S.S.D, J.F.A. and N.C.R performed experiments. C.W. and S.S.D developed the algorithm and performed computational analysis. C.W. and S.S.D wrote the manuscript.

## Methods

### Embryo isolation and cell picking

Embryos were gently flushed out of the infundibulum of E2.5 pregnant mice using warm M2 medium. Embryos were then manipulated in 4-ring IVF dishes coated with RNase-free BSA because, once dissociated, cells stick to plastic. Embryos were washed in PBS-0 and in Tyrode’s acid to remove the zona pellucida, then placed in a 1/3 dilution of TrypLE Select Gibco A12177-01 (stock solution is referred by Gibco as 10x concentrated) and placed on the warm plate for 2 minutes. We then used glass capillaries of different diameters to dissociate the embryo into 2-3 clusters. Cells were then progressively extracted from each cluster, one after the other, using glass capillaries. Every single cell that is released from the clusters is immediately placed into a well of a 384-well plate containing lysis buffer.

### Single-cell 5hmC sequencing (scAba-Seq)

Single cells isolated from 8-cell mouse embryos were deposited into 384-well plates and the scAba-Seq protocol was performed using the Nanodrop II liquid-handing robot. Briefly, after protease treatment to strip off chromatin, 5hmC sites in the genome were glucosylated using T4-Phage β-glucosyltransferase. Next, AbaSI was added to the reaction mixture that recognizes glucosylated sites and introduces double-stranded breaks with 3’ overhangs 11-13 nucleotides downstream of the recognition site. The fragmented genomic DNA molecules were ligated to double-stranded adapters containing a cell barcode, 5’ Illumina adapter, and T7 promoter. The ligated molecules were amplified by *in vitro* transcription and then used to prepare Illumina libraries. A detailed protocol can be found in Ref. 14.

### Modeling SCE events as a Poisson process

The 5hmC data was discretized into 2 Mb bins and all SCE transitions in the 8-cell mouse embryos were identified manually. A specific SCE transition on chromosome 14 was found at the same genomic position in all embryos due to a misorientation of the reference genome (mm10), consistent with previous reports^18,21^. The stochastic nature of SCE events is modeled as a Poisson process. In using a Poisson process to model SCE events, we assume that all SCE events occur independently and at a constant rate. The probability of observing *x* SCE transitions in one cell cycle is given by:

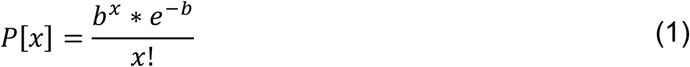

where *b* is the average number of SCE transitions per chromosome per cell division. Further, to build a probabilistic framework to reconstruct cellular lineages, we define the following parameters: (1) *r* is the probability that an original strand is inherited by a particular daughter cell, which is equal to ½ for randomly segregating DNA strands; (2) *k*_*ij*_ is the genomic length fraction of the *j*^th^ segment (1 ≤ *j* ≤ *l* + 1, where *l* is the number of SCE transitions) of the original DNA strand that is observed in cell *i*; and (3) *N* is the number of unique positions where SCE events can occur.

### scPECLR

The first step is to use the numbers of observed SCE events to estimate *b* using maximum likelihood estimation (MLE). Thereafter, Original Strand Segregation (OSS) analysis is used to separate the cells into two groups, reducing the number of cell divisions to be reconstructed from *n* to *n* − 1. Next, within each subtree, we calculate the probability of observing a SCE pattern of a chromosome given a tree topology. For example, for the most frequently occurring pattern of one SCE event shared between two cells (see example in Figure 2-I), the probability of observing it in Tree A is given by the product of the probability of having no SCE events in the first cell division and the probability of having one SCE event in the second cell division

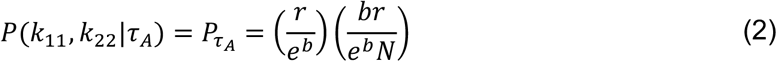

Similarly, the probability of observing this pattern in Tree B is given by the product of the probability of having one SCE event in the first cell division, and no SCE events within the original DNA strands in both cells in the second cell division

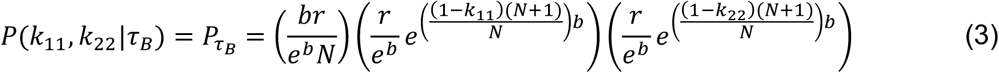

which leads to

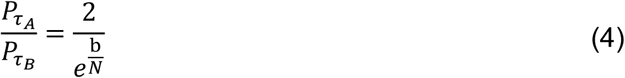

Detailed analytical expressions for the probability of observing different SCE patters are provided below in “Analytical expressions for the probability of observing the three most common SCE patterns”.

Subsequently, we assume that the SCE patterns on each chromosome are independent and compute the overall probability of observing SCE events over the whole genome for each tree topology. Moreover, as a 4-cell subtree has only three distinct topologies we get

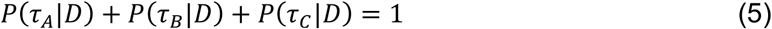

where *D* represents the genome-wide SCE patterns in all cells of the embryo. Rearrangement gives us the probability of observing different tree topologies given the SCE patterns over the whole genome

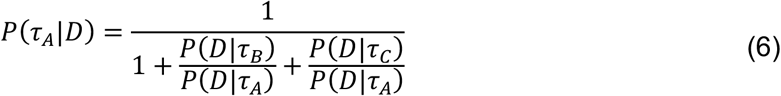

Finally, the probability of observing the topology of a particular 8-cell tree is a product of the probabilities of the two corresponding 4-cell subtrees.

After the probabilities of all tree topologies are estimated, scPECLR assigns the topology with the highest probability as the predicted tree. Then, starting with this predicted tree, *b* values specific to each cell division are estimated. A second iteration with cell division-specific *b* values is then performed to obtain a new predicted tree. If the new predicted tree is not the same tree as that inferred in the first iteration, another iteration is performed starting from the predicted tree in the current iteration. This iterative process is carried out till the predicted tree is the same as that obtained in the previous iteration or until 10 iterations have been performed. In all *in vivo* mouse embryos and almost all simulated embryos, the predicted tree converges by the 3^rd^ iteration.

### Analytical expressions for the probability of observing the three most common SCE patterns

#### Case I

The most common SCE pattern that we observed in mouse embryos is one SCE transition shared between two cells (cells 1 and 2 in Figure 2-I and Supplementary Figure 1). This pattern alone cannot discriminate between sister (Tree A) or cousin (Trees B and C) cell configurations as all three topologies are consistent with the SCE pattern. Therefore, we developed a model to rigorously determine the probability of observing any SCE pattern given a tree topology. For Tree A, the probability of observing one shared SCE transition is given by the product of the probability of having no SCE events in the first cell division and the probability of having one SCE event in the second cell division. Further, there is a ^1^/_*N*_ chance that the observed SCE event occurs at a specific discretized genomic position. The probability that the original DNA strand is inherited by the mother of cells 1 and 2 is *r*, and the probability of inheriting the observed SCE pattern between cells 1 and 2 is given by *r*.

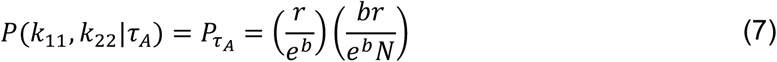

Similarly, for Tree B,

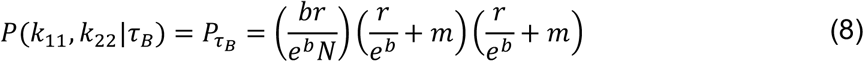

Here, *m* represents the probability that the SCE events during the second cell division occur within newly synthesized DNA strands that contain undetectable levels of 5hmC. To estimate *m* on the left branch of the lineage tree that gives rise to cells 1 and 3, we can show that

Probability of 1 undetectable SCE 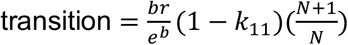

Probability of 2 undetectable SCE 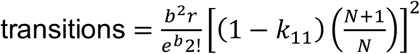

Probability of *n* undetectable SCE 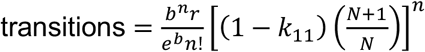

Therefore, *m* is given by

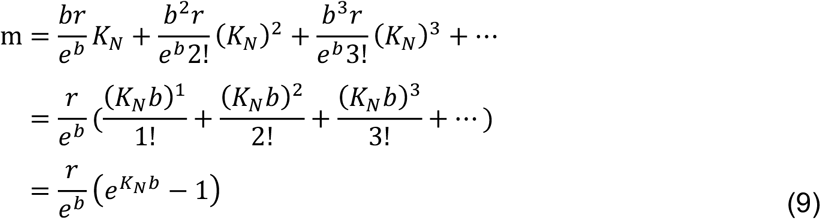

where 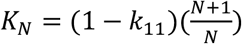.

Thus, (8) becomes

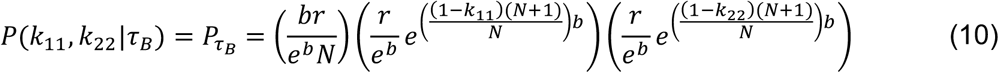

Further, it is trivial to show that the probability of observing the SCE pattern given Tree B or C is equal, that is

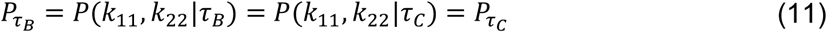

Therefore, the ratio of the probability of cells 1 and 2 being sisters (Tree A) *vs*. cousins (Trees B or C) is given by

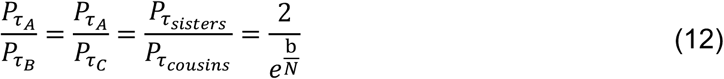

Note that the probability ratio is a function of only the SCE rate and the number of bins, and is not dependent on the location of the SCE event in this case.

#### Case II

The second most common SCE pattern is the observation of two SCE transitions that are shared between two cells (Figure 2-III and Supplementary Figures 1 and 2). For the original DNA strand to be observed in only two cells, SCE transitions must occur in the same cell cycle. Thus, the probability of observing this SCE pattern in Tree A is given by

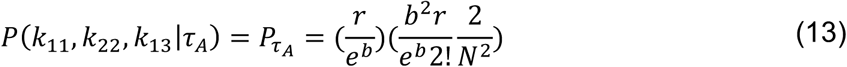

The first term is the probability that no SCE event occurs in the first cell division, and the second term is the probability of having two SCE transitions during the second cell division.

Similarly, for Tree B

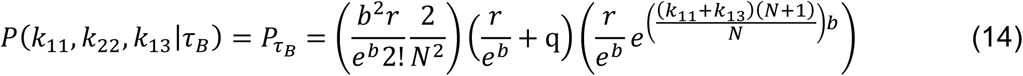

where *q* is the probability that undetectable SCE events occur within the 5hmC-depleted genomic region between *k*_11_ and *k*_13_, whose length is equal to *k*_22_. Note that the observed SCE pattern is possible for an even number of SCE events occurring within this region. To estimate *q*, we can show that

Probability of 2 undetectable SCE 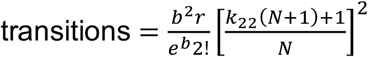

Probability of 4 undetectable SCE 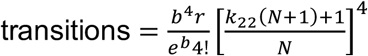

Probability of *n* undetectable SCE 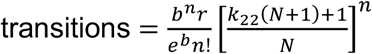

Thus, *q* is given by

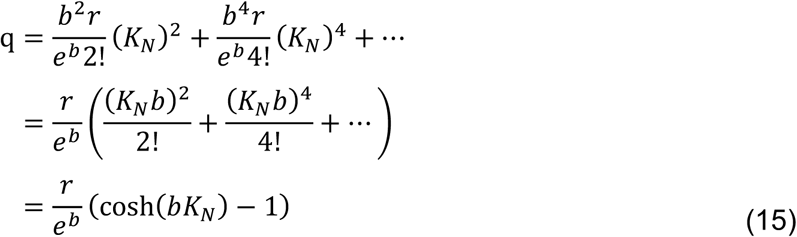

where 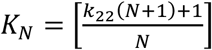

Therefore, (14) becomes

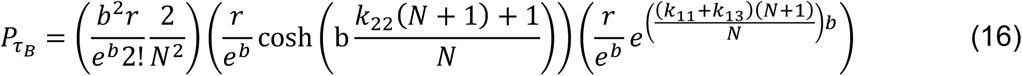

and the ratio of the probability of cells 1 and 2 being sisters (Tree A) *vs*. cousins (Trees B or C) is given by

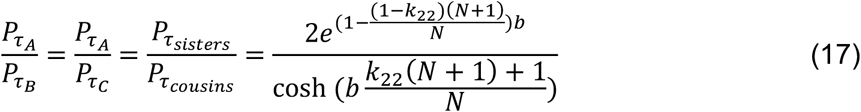

In this case, the probability ratio is a function of the genomic location of the SCE events, in addition to the SCE rate and the number of bins.

#### Case III

Another common but more complicated SCE pattern occurs when an original DNA strand is shared between three cells (Figure 2-III). Intuitively, Tree B with cells 1 and 3 as sisters is the least likely configuration as it requires one additional SCE transition compared to the other two trees. The probability of observing this SCE pattern in Trees A and C are given by

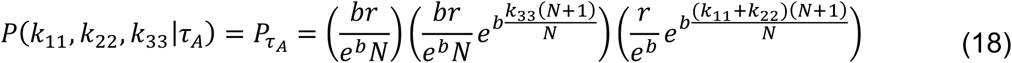

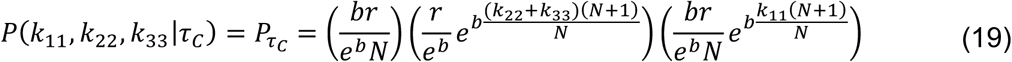

In (18), the first term accounts for one SCE event between *k*_22_ and *k*_33_. The second term includes one SCE event between *k*_11_and *k*_22_ and undetectable SCE events within the right-most genomic region, whose length is equal to *k*_33_. The third term accounts for no SCE event within *k*_33_ and undetectable SCE events within the left region, whose length is equal to (*k*_11_ + *k*_22_). Similarly, in (19), the first term accounts for one SCE event between *k*_11_ and *k*_22_. The second term includes no SCE events within *k*_11_ and undetectable SCE events within the rest of the chromosome, equivalent in length to (*k*_22_ + *k*_33_). The third term includes one SCE event between *k*_22_ and *k*_33_ and undetectable SCE events within the left-most genomic region. Note that Trees A and C are mirror images of each other and the probability of observing this SCE pattern is equal for these two tree configurations. For Tree B,

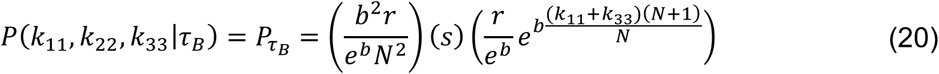

The first term is for two SCE events in the first cell division. The second term accounts for an odd number of undetectable SCE transitions within the genomic region between *k*_11_ and *k*_33_, such that both cells 1 and 3 contain parts of the original DNA strand. The third term includes undetectable SCE events within both left and right genomic regions, whose combined length is (*k*_11_ + *k*_33_). Further, *s* is given by

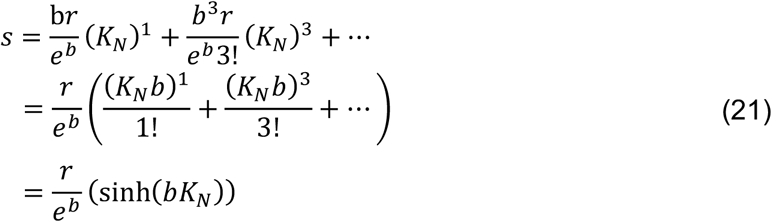

where 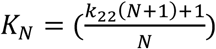.

Therefore, (20) becomes

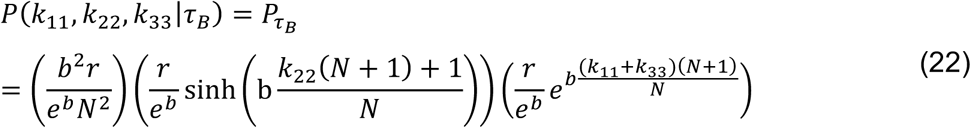

and the ratio of the probability of Tree A *vs*. B is given by

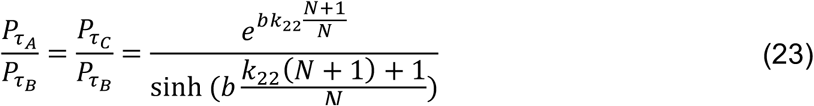

Consistent with our intuition, Tree B is less likely than the other two tree topologies, and depending on the values of *N, b*, and *k*_22_, Tree B can be anywhere between 2 to 100 times less likely (Figure 2-III).

The approach described above can be applied to any SCE pattern. The probability of observing different SCE patterns are estimated for all chromosomes. Next, we assume that each chromosome strand is independent and compute the overall probability of observing the SCE patterns over the whole genome (*D*) for each Tree *i* (*τ*_*i*_). To determine the most likely tree, we compute and compare *P*(*τ*_*A*_|*D*), *P*(*τ*_*B*_|*D*), and *P*(*τ*_*C*_|*D*) using Bayes’ theorem

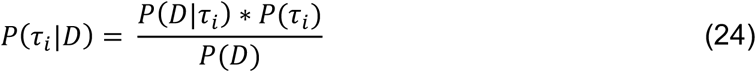

where *P*(*τ*_*i*_) and *P*(*D*) are the probabilities of observing Tree *i* and the genome-wide SCE pattern data, respectively. *P*(*τ*_*i*_) reflects prior belief of the likelihood that Tree *i* is the correct topology. As there are 3 possible topologies for any 4-cell tree, we get

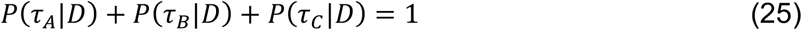

Further, the ratio of the probability of observing Tree *i vs*. Tree *j* is given by

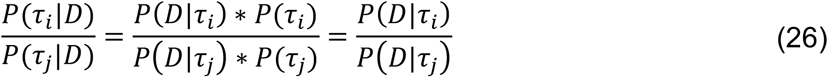

where Tree *i or j* is either Tree *A, B*, or *C*. The prior probabilities *P*(*τ*_*i*_) are assumed to be equal to one another, a common practice in Bayesian analysis^31^. After rearrangement, we get

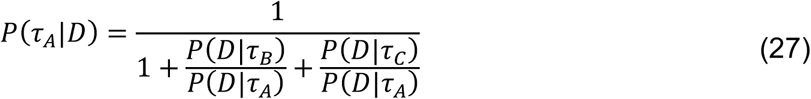

Similarly, the probability of all tree topologies can be calculated. Finally, the probability of a particular 8-cell tree is given by the product of the probabilities of the two corresponding 4-cell subtrees.

### Simulating stand-specific 5hmC distributions

To validate the analytical expressions for the probability of observing different SCE patterns in Figures 2-II and 2-III, we simulated 8-cell trees where the occurrence of SCE events were modeled as a Poisson process with *b* = 0.3 and chromosome strands were assumed to segregate randomly (*r* = 0.5). Simulations were performed on chromosome 1 (*N* = 97 for 2 Mb bins). These simulations were then used to estimate the probability of observing Tree A *vs*. Tree B as a function of the position of the SCE event.

To test the accuracy of scPECLR in predicting lineage trees in Figure 3-II, 8- or 16-cell embryos with 19 or 38 chromosomes were simulated as described above. All bins in the original DNA strands were hydroxymethylated whereas all subsequently synthesized DNA strands contained no 5hmC, mimicking *in vivo* experimental observations. 5,000 simulated trees were generated for each condition shown in Figure 3-II and inputted into scPECLR to estimate the percentage of trees that are accurately predicted by the algorithm. For 16-cell trees, we also estimated the prediction accuracy of 2-, 4- and 8-cell subtrees within the full tree. For 4-cell embryos, as OSS accurately separates the four cells into two groups of two cells each, the lineage reconstruction problem becomes deterministic and the trees are predicted with 100% accuracy.

### Consensus tree analysis

This analysis was performed on 16-cell trees to identify parts of the lineage tree that can be predicted with high confidence. The two 8-cell subtrees obtained from OSS are treated independently. The first step is to use a desired relative threshold (RT) to identify all trees that have probabilities within a threshold level of the highest probability tree and include such trees for downstream analysis. All included trees are subsequently weighed equally. The second step is to examine the 4-cell subtrees of each included tree. If all trees consistently predict the same 4-cell subtree, the consensus tree includes the 4-cell subtree. This is true for most datasets as scPECLR largely predicts the 4-cell subtrees accurately in 16-cell trees (Figure 3-III). When disagreement arises, if the percentage of included trees that have the same 4-cell subtree exceeds a threshold (48), ranging from 0.55 to 1.0, the consensus tree includes the 4-cell subtree, and tree topologies that conflict with this 4-cell subtree are excluded from further analysis. If the percentage is below *t*8, the consensus tree does not include the exact 4-cell subtree but instead attempts to identify as many pairs of cells as possible that appear in different 4-cell subtrees of all included trees, and the consensus analysis terminates. After the 4-cell subtrees are determined, the topology predicted within each of these subtrees is then considered. Again, if all of the remaining trees predict the same topology or if the percentage of remaining trees that predict a consistent topology exceeds a threshold (*t*4), ranging from 0.55 to 1.0, the consensus tree also includes that topology. Otherwise, it does not predict a specific topology within the 4-cell subtree but attempts to identify one cousin pair that appears in the 4-cell topology.

The consensus tree has different levels of specificity, ranging from predicting a full 16-cell tree, where the relationships between all cells are exact, to predicting only two 8-cell subtrees. In general, each consensus tree is constrained to contain a certain number of tree topologies, which provides information about how specific each consensus tree is. For example, in Figure 3-IV, the consensus tree contains six possible topologies, as there are two topologies arising from uncertainty in the subtree containing cells 5-8 and three topologies arising from uncertainty in the subtree containing cells 13-16. The lower the number of topologies contained within the consensus tree, the more specific and informative it is.

There are three parameters in the consensus tree analysis: RT, *t*8, and *t*4. RT has the largest influence on the structure of the consensus tree, while varying *t*8 and *t*4 leaves the consensus tree largely unchanged (Figures 3-V, 3-VI and Supplementary Figure 4) (Note: In Figure 3-V, 48 and *t*4 are kept constant at 0.75 and 1, respectively). When the RT increases, the consensus tree becomes more specific but suffers from a higher false discovery rate (FDR). In contrast, although the effects are small, increasing *t*8 and *t*4 leads to a very modest decrease in the specificity of the consensus tree and reduction in FDR. Thus, using different parameter values allows us to tune the competing goals of specificity and accuracy of the consensus tree. In fact, for a specific FDR, there is an optimal set of parameters that gives the most specific consensus tree for a dataset. We performed a consensus tree analysis on the dataset in Figure 3-II (solid blue lines), with different combinations of RT ranging from 0.05 to 0.50, and *t*8 and *t*4 ranging from 0.55 to 1.0. Each parameter set provides a consensus tree with a different level of specificity, measured by the median number of trees contained in the consensus tree, and the FDR. For any level of FDR tolerated, there is at least one parameter combination that yields the lowest median number of trees. For example, when *b* = 0.3 and the FDR is chosen to be 30%, the optimal parameter set has RT, *t*8, and *t*4 as 0.05, 0.75, and 1, respectively, yielding the median number of trees contained within the consensus tree to be 36. Thus, for any dataset, the rate of SCE events can be estimated using MLE, and with a user-selected FDR, an optimal parameter set can be estimated to give the most specific consensus tree.

Consensus tree analysis improves the accuracy of lineage prediction in all scenarios. When the SCE rate is low (*b* = 0.1) and the iterative prediction alone performs poorly for 16-cell trees, an error rate of greater than 99% in the iterative prediction decreases to a FDR between 30-75%. When the iterative prediction alone performs moderately (*b* = 0.5), an error rate of ∼60% improves to a FDR between 10-45% (Figures 3-II and 3-V). Lastly, when the iterative prediction alone performs well (*b* = 1.0), an error rate of ∼25% decreases to a FDR between 5-20% (Figures 3-II and 3-V). When *b* = 1.0, there are only 1 to 2 median topologies contained in each consensus tree, indicating that the consensus analysis increases the accuracy of the prediction without compromising its specificity. This result shows that scPECLR and the consensus tree analysis provides a significant amount of lineage information with reasonable accuracy for 16-cell trees (Figures 3-V and 3-VI).

### scPECLR is robust to initial estimates of the SCE rate and to varying SCE rates at each cell division

We explored the robustness of scPECLR to initial estimates of the SCE rate by simulating strand-specific 5hmC data in 8-cell trees with a constant SCE rate (*b* = 0.3). We then used different values of SCE rates – ranging from 0.1 to 2.0 – in scPECLR to predict the lineage tree (instead of estimating the SCE rate from the observed SCE pattern using MLE). We found that the percentage of trees that were accurately predicted did not change over the range of SCE rates, suggesting that scPECLR is robust to uncertainty in SCE rate estimation (Supplementary Figures 5-I and 5-II).

As the 8-cell mouse embryos have varying rates of SCE events across cell divisions, we explored the robustness of scPECLR when the rates are different for each cell division. Because prediction accuracy of scPECLR is dependent on the rate of SCE events, in this analysis, we fixed the combined SCE rate (*B*) over 3 (or 4) cell divisions, but allowed individual cell divisions to have different rates. For 8-cell trees, the model is largely robust against varying rates of SCE events across cell divisions, with higher *B* and larger number of chromosomes resulting in better prediction accuracy (Supplementary Figure 5-III). For example, when the SCE rates are low for the first and second cell division (*b*_1_ and *b*_3_) and high for the third cell division (*b*_3_), similar to the experimental observation in 8-cell mouse embryos, scPECLR predicts the lineage tree with very high accuracy (Supplementary Figure 5-III, H3). One case where the prediction accuracy drops modestly is when the SCE rates of the first and third cell divisions (*b*_1_ and *b*_3_) are low and the SCE rate of the second cell division (*b*_3_) is high (Supplementary Figure 5-III, H2). In this case, the data has a large number of SCE events that are shared between cousin cells. As the SCE rate at each cell division is assumed constant during the first iteration of scPECLR, the algorithm predicts that cells sharing more SCE events are more likely to be sisters. This misidentification results in a large percentage of simulations not predicting the true tree after the first iteration. However, the prediction improves significantly after a few iterations because starting from the second iteration, the model accounts for different SCE rates at each cell division. Consequently, the varying SCE rates at each cell division has minimal impact on the accuracy of 8-cell tree prediction.

For 16-cell trees, there are a few cases where the prediction accuracy is worse than when the rates are uniformly distributed; these include situations where *b*_4_ is low (Supplementary Figure 5-IV, H2, H3, H13, H23, and L4). In these cases, the prediction accuracy is lower because scPECLR inaccurately infers a pair of cousin or second cousin cells as sister cells due to a large number of SCE events shared between such pairs. In contrast, cases with high *b*_4_ values result in better prediction accuracy because scPECLR correctly identifies sister cell pairs (Supplementary Figure 5-IV, H4, H14, H24, and H34). Finally, scPECLR also performs well when *b*_2_ and *b*_3_ are low as it does not misidentify cousin or second cousin pairs as sister pairs. These results suggest that in addition to the combined SCE rate, how the individual SCE rates are distributed over each cell division impacts the accuracy of reconstructing 16-cell trees.

### Binomial test to identify non-random DNA segregation

To test the segregation pattern of DNA strands at the 4-cell stage, the 5hmC profile of 8-cell mouse embryos were combined using the lineages predicted by scPECLR to obtain the distribution of 5hmC on the original DNA strands at the 4-cell stage. Original DNA strands that had not undergone any SCE events at the 4-cell stage were considered in this analysis. Due to low SCE rates during the first and second cell divisions in the embryos, a majority of the original DNA strands had not undergone SCE events at the 4-cell stage. A binomial two-tailed test was conducted in *R* with a null hypothesis of random segregation (*π* = 0.5) and an alternative hypothesis of non-random segregation (*π* ≠ 0.5). A pair of sister cells were considered to display statistically significant non-random DNA segregation for p-values lower than 0.05.

### scPECLR implementation in MATLAB

scPECLR was implemented in MATLAB to perform iterative probabilistic reconstruction of lineage trees. The script first uses single-cell strand-specific 5hmC data to perform OSS analysis to eliminate a majority of tree topologies. Next, it calculates the SCE rate and estimates the probabilities of all tree topologies given the genome-wide SCE pattern to predict the tree with the highest probability. Using this predicted tree, the program estimates the SCE rate for each cell division and re-calculates the probabilities of all tree topologies. The program performs iterations until the predicted tree does not change or until 10 iterations are reached. The scripts implementing scPECLR in MATLAB, along with test files, are provided as Supplementary Information.

### Data and software availability

Accession code GEO: GSE131678. scPECLR was implemented in MATLAB. Custom codes and test files are provided with this manuscript.

