## Supplementary Information for "A probabilistic framework for cellular lineage reconstruction using single-cell 5-hydroxymethylcytosine sequencing"

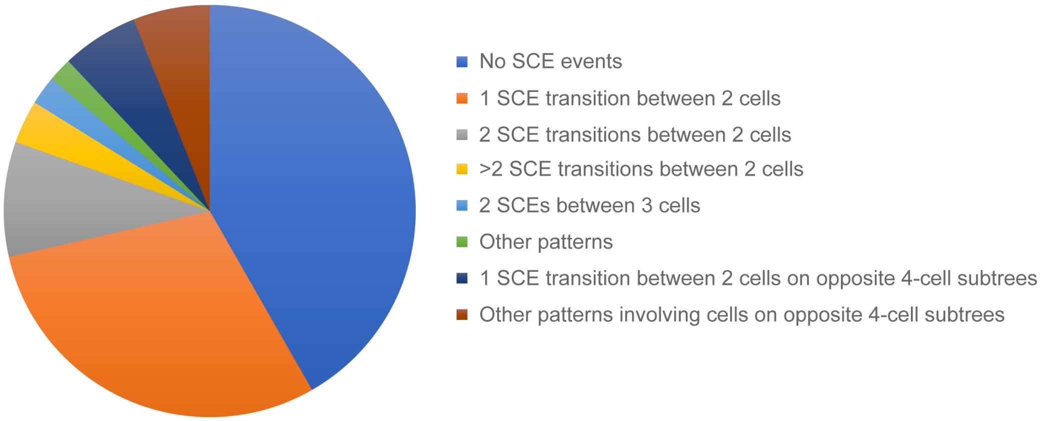


**Supplementary Figure 1 |** Distribution of SCE patterns in 8-cell mouse embryos. Approximately 33% of the original DNA strands display one SCE transition that is shared between two cells within the same 4-cell subtree (orange). In addition to this most frequently observed pattern, a large diversity of other SCE patterns are observed in 8-cell mouse embryos. All observed SCE patterns are used to probabilistically reconstruct cellular lineages in scPECLR. Approximately 40% of the original DNA strands do not undergo SCE events in the first three cell divisions up to the 8-cell stage of mouse embryogenesis.

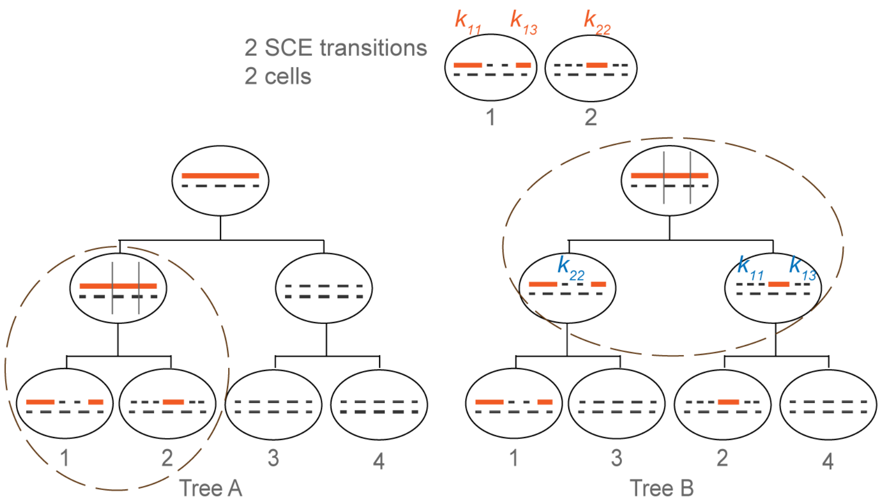


**Supplementary Figure 2 |** The more complex pattern of two SCE transitions shared between two cells increasingly favors the sister tree to the cousin tree arrangement. Schematic showing the SCE events that are necessary for two cells that share two SCE transitions to be sister (Tree A) or cousin cells (Tree B). Mathematically, the original DNA strand undergoes the same number of SCE transitions in both tree topologies and the probability of observing the SCE event shown within the dotted circles is identical for Trees A and B. Further, in Tree A, the cell division that gives rise to cells 3 and 4 is unconstrained in the number of SCE events that can take place. In contrast, while any number of SCE events can occur within the $k_{11}$ and $k_{13}$ genomic regions in Tree B, the $k_{22}$ region is constrained to have an even number of SCE events, thereby reducing the likelihood to observing Tree B compared to Tree A.

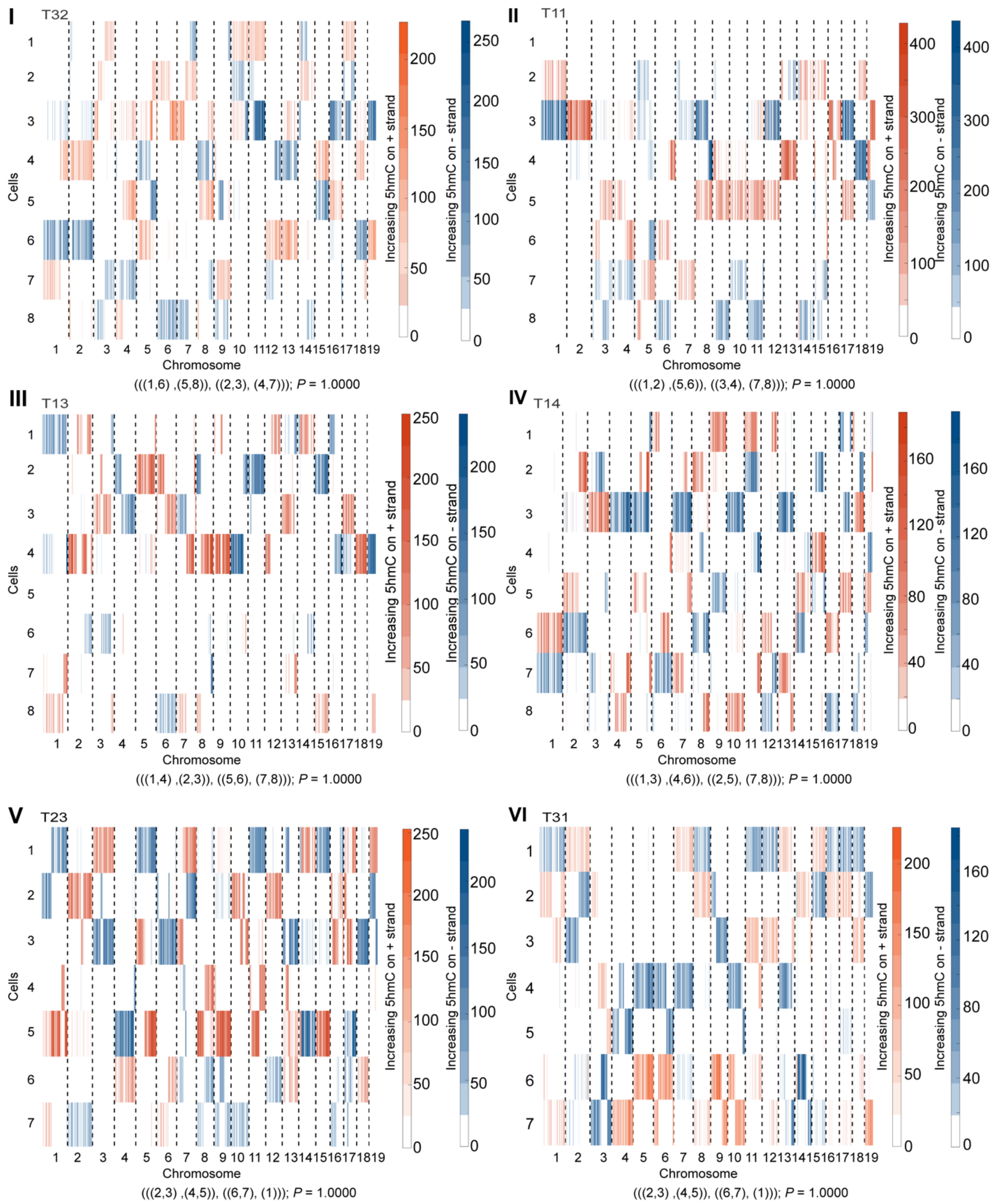


**Supplementary Figure 3 |** Reconstructing lineage trees for preimplantation mouse embryos using scPECLR. (I-VI) The genome-wide strand-specific 5hmC distribution of 4 8-cell and 2 7-cell mouse embryos are shown. The 7-cell mouse embryos contain one blastomere from the 4-cell stage of embryogenesis that had not divided at the time the embryos were isolated. The mosaic pattern of 5hmC and the SCE events can be used to reconstruct the cellular lineages using scPECLR. The predicted lineage tree and the probability of observing this topology is indicated below each panel. Reconstructing the 7-cell embryos T23 and T31 shows that scPECLR can be applied to non-symmetric trees. Note that for two 8-cell mouse embryos we were not able to successfully sequence the 5hmC of one cell in each embryo (cell 1 in the embryo T11 and cell 5 in the embryo T13). By assuming that the original DNA strands that were not observed in any of the remaining 7 cells must have been present in the cell that failed to sequence, we were able to successfully predict the 8-cell lineage tree. These results suggest that scPECLR can also be used in cases where there is a limited amount of missing 5hmC sequencing data.

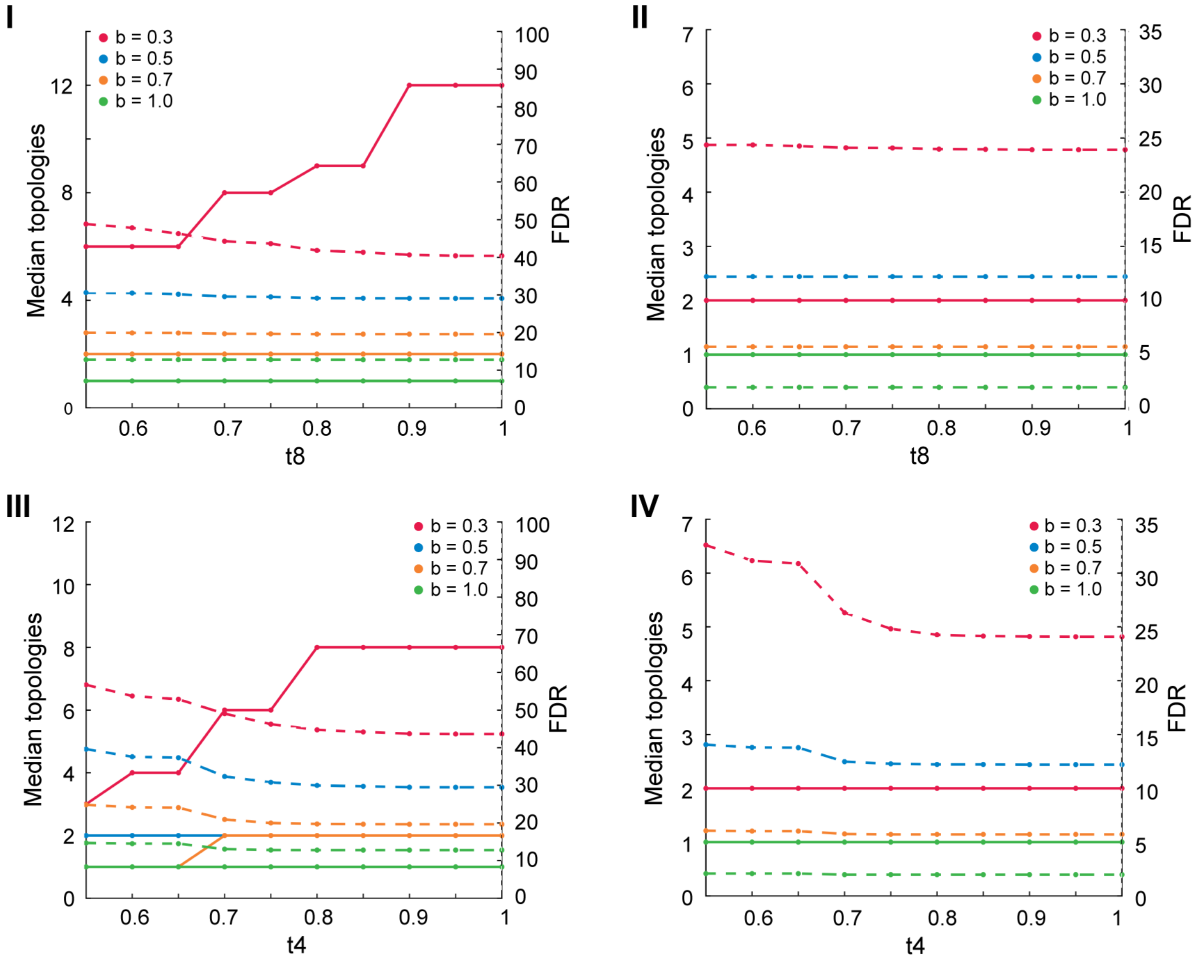


**Supplementary Figure 4 |** Parameters $t8$ and $t4$ have minor impact on the consensus tree analysis. Panels show representative examples of how the median number of topologies of the consensus tree and the FDR varies with $t8$ and $t4$. These plots are shown for (I) $RT = 0.25$ and $t4 = 1$ for 16-cell trees with 19 chromosomes; (II) $RT = 0.25$ and $t4 = 1$ for 16-cell trees with 38 chromosomes; (III) $RT = 0.25$ and $t8 = 0.75$ for 16-cell trees with 19 chromosomes; and (IV) $RT = 0.25$ and $t8 = 0.75$ for 16-cell trees with 38 chromosomes. Solid lines indicate the median number of topologies contained in the consensus tree on the left axis, and the dotted lines indicate the FDR on the right dotted axis. Varying $t8$ and $t4$ across the entire range of values shows that it does not have a significant impact on the median number of topologies contained in the consensus tree or the FDR. Note that in panel (I), the blue solid line is covered by the yellow solid line as they have the same number of median topologies in all cases. Similarly, in panels (II) and (IV), both the blue and yellow solid lines are covered by the green solid line.

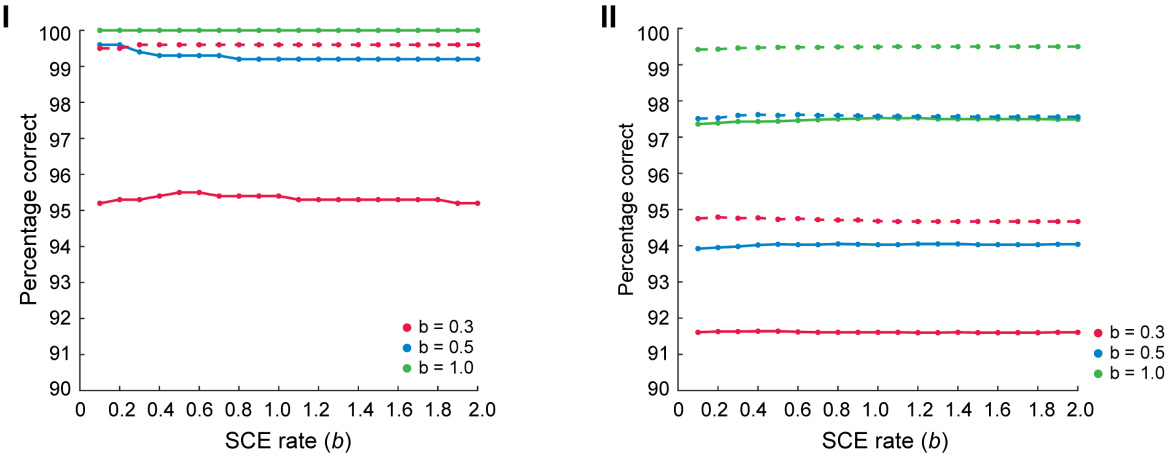


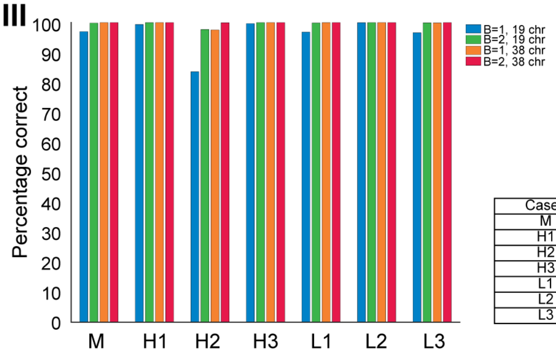


| Cases | $b_1$ | $b_2$ | $b_3$ |
|---|---|---|---|
| M | 0.33 / 0.67 | 0.33 / 0.67 | 0.33 / 0.67 |
| H1 | 0.6 / 1.2 | 0.2 / 0.4 | 0.2 / 0.4 |
| H2 | 0.2 / 0.4 | 0.6 / 1.2 | 0.2 / 0.4 |
| H3 | 0.2 / 0.4 | 0.2 / 0.4 | 0.6 / 1.2 |
| L1 | 0.2 / 0.4 | 0.4 / 0.8 | 0.4 / 0.8 |
| L2 | 0.4 / 0.8 | 0.2 / 0.4 | 0.4 / 0.8 |
| L3 | 0.4 / 0.8 | 0.4 / 0.8 | 0.2 / 0.4 |

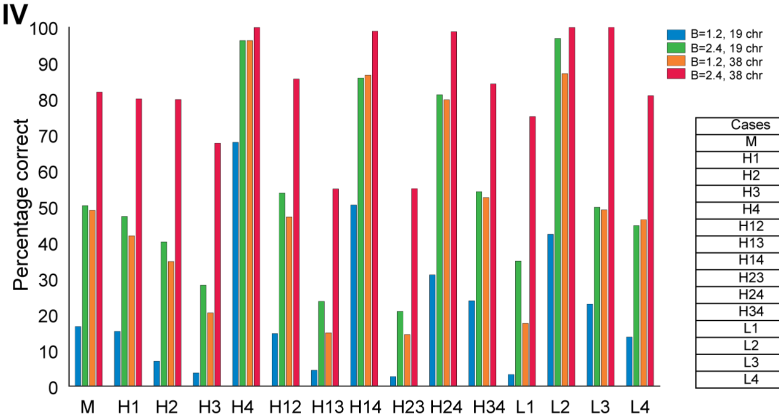


| Cases | $b_1$ | $b_2$ | $b_3$ | $b_4$ |
|---|---|---|---|---|
| M | 0.3 / 0.6 | 0.3 / 0.6 | 0.3 / 0.6 | 0.3 / 0.6 |
| H1 | 0.6 / 1.2 | 0.2 / 0.4 | 0.2 / 0.4 | 0.2 / 0.4 |
| H2 | 0.2 / 0.4 | 0.6 / 1.2 | 0.2 / 0.4 | 0.2 / 0.4 |
| H3 | 0.2 / 0.4 | 0.2 / 0.4 | 0.6 / 1.2 | 0.2 / 0.4 |
| H4 | 0.2 / 0.4 | 0.2 / 0.4 | 0.3 / 0.6 | 0.6 / 1.2 |
| H12 | 0.4 / 0.8 | 0.4 / 0.8 | 0.2 / 0.4 | 0.2 / 0.4 |
| H13 | 0.4 / 0.8 | 0.2 / 0.4 | 0.4 / 0.8 | 0.2 / 0.4 |
| H14 | 0.4 / 0.8 | 0.2 / 0.4 | 0.2 / 0.4 | 0.4 / 0.8 |
| H23 | 0.2 / 0.4 | 0.4 / 0.8 | 0.4 / 0.8 | 0.2 / 0.4 |
| H24 | 0.2 / 0.4 | 0.4 / 0.8 | 0.2 / 0.4 | 0.4 / 0.8 |
| H34 | 0.2 / 0.4 | 0.2 / 0.4 | 0.4 / 0.8 | 0.4 / 0.8 |
| L1 | 0.17 / 0.34 | 0.34 / 0.69 | 0.34 / 0.69 | 0.34 / 0.69 |
| L2 | 0.34 / 0.69 | 0.17 / 0.34 | 0.34 / 0.69 | 0.34 / 0.69 |
| L3 | 0.34 / 0.69 | 0.34 / 0.69 | 0.17 / 0.34 | 0.34 / 0.69 |
| L4 | 0.34 / 0.69 | 0.34 / 0.69 | 0.34 / 0.69 | 0.17 / 0.34 |

**Supplementary Figure 5 |** scPECLR is robust to initial estimates of the SCE rate and to varying SCE rates at each cell division. The sensitivity of the prediction accuracy of scPECLR to initial estimates of the SCE rate was tested for (I) 8-cell and (II) 16-cell trees. Trees were simulated with a constant SCE rate of $b = 0.3$ (red), $b = 0.5$ (blue), $b = 1.0$ (green). To test the robustness of the algorithm, instead of estimating the SCE rate from the data, values ranging from 0.1 to 2.0 were used during the first iteration of scPECLR to predict the tree. We found that the percentage of trees that were accurately predicted was robust across the range of SCE rates for cells containing both 19 (solid lines) and 38 chromosomes (dotted lines). Note that in panel (I), the dotted lines corresponding to $b = 0.5$ and $b = 1.0$ are hidden behind the solid line corresponding to $b = 1.0$, as they all have 100% prediction accuracy for all SCE rates. These results are based on 1000 simulated trees for each condition. (III) Panel shows the percentage of 8-cell trees (with cells containing 19 or 38 chromosomes) that are accurately predicted for varying SCE rates over the 3 cell divisions. To systematically compare the prediction accuracy of scPECLR, the combined SCE rate ($B = b_1 + b_2 + b_3$) is held constant over all cell divisions. The results show that scPECLR can accurately predict the lineage of 8-cell trees when the SCE rates vary with each cell division. (IV) Panel shows the percentage of 16-cell trees (with cells containing 19 or 38 chromosomes) that are accurately predicted for varying SCE rates over the 4 cell divisions. $M$ denotes cases where the SCE rate is constant over all cell divisions. $H_i$ (and $L_i$) denotes cases where the SCE rate is higher (or lower) in the $i^{th}$ cell division, and $H_{ij}$ denotes cases where the SCE rate is higher in the $i^{th}$ and $j^{th}$ cell division over the other two cell divisions. These results are based on 5000 simulated trees for each condition.
