## Supplementary material for "A probabilistic framework for cellular lineage reconstruction using single-cell 5-hydroxymethylcytosine sequencing": Code (scPECLR): Determine_Break_Num.m

```
%% Determine the number of breaks
% input forward and reverse strands (with chr and bin# as first two columns) 
% determine the break number for each chromosome and b_observed fit with MLE
% NOTE THAT THE SAME BREAK IN THE FIRST CELL DIVISION WILL BE COUNTED TWICE
% BUT THAT SHOULD NOT AFFECT B_OBS MUCH
function [brk_num, b_obs] = Determine_Break_Num(cell_f,cell_r)
    chr_num = max(cell_f(:,1)); 
    cell_num = size(cell_f,2) - 2; 
    cd_num = log2(cell_num); %cell division number 
    brk_num = zeros(chr_num,3); % 1st col = chr#, 2nd/3rd col = #break on +,- 
    for i = 1:chr_num
        ii1 = find(cell_f(:,1) == i); % find the rows
        f_chr = cell_f(ii1,3:end);
        r_chr = cell_r(ii1,3:end);
        siz = size(f_chr,1);
        pos_count = 0;
        neg_count = 0; 
        for j = 1:cell_num
            f_holder = f_chr(1,j);
            r_holder = r_chr(1,j);
            for k = 2:siz
                if f_chr(k,j) ~= f_holder
                   pos_count = pos_count + 1;
                   f_holder = f_chr(k,j);
                end
                if r_chr(k,j) ~= r_holder
                   neg_count = neg_count + 1;
                   r_holder = r_chr(k,j);
                end           
            end
        end
        brk_num(i,1) = i;
        brk_num(i,2) = pos_count/2; % divide by 2 to get break num
        brk_num(i,3) = neg_count/2;
    end
    data = [brk_num(:,2); brk_num(:,3)]; %all break num
    b_full = poissfit(data); %get value of b fit with MLE
    b_obs = b_full/cd_num; % divide to get b per cell div 
end
```
