## Supplementary material for "A probabilistic framework for cellular lineage reconstruction using single-cell 5-hydroxymethylcytosine sequencing": Code (scPECLR): OSS_16.m

```
function [Ptotal, left_side, right_side, totbin] = OSS_16(SixteenCell_f,SixteenCell_r)
% generate tree matrix Ptotal 8 cells
% input forward and minus strand in 1/0 format with no overlap.
% output relevant Ptotal after OSS, cells in left side, cells in right side, totbin  
cell_num = 16;
v = 1:cell_num;
% need to seperate the two sets; find the start and stop of each chromosome
Coord = SixteenCell_f(:,1:2);
start(1) = 1;
for i = 2:1:length(Coord(:,1))
    if Coord(i,1) ~= Coord(i-1,1)
        stop(Coord(i-1,1)) = i-1;
        start(Coord(i,1))  = i;
    end
end
stop(Coord(i,1)) = i;
totbin = stop - start + 1;
left_side = zeros(cell_num/2,1); %cells on the left side
right_side = zeros(cell_num/2,1); %cells on the right side
left_side(1,1) = 1; %arbitrarily places cell 1 on the left side  
% find the bins where cells 1 has the orignal plus and minus strand
TF_left_plus = SixteenCell_f(:,left_side(1,1)+2) == 1; 
TF_left_minus = SixteenCell_r(:,left_side(1,1)+2) == 1; 
TF_left = TF_left_plus | TF_left_minus;
% if any other cell has any of these bins -> on the right
cells_on_right_bin = SixteenCell_f(TF_left,3:end) | SixteenCell_r(TF_left,3:end);
if size(cells_on_right_bin,1) == 1 % in case there is only 1 bin and we cannot use any
    cells_on_right = cells_on_right_bin;
else
    cells_on_right = any(cells_on_right_bin);
end
% not cell 1 - remove that
cells_on_right(1,1) = 0;
cells_right = find(cells_on_right == 1);
for i = 1:sum(cells_on_right)
    right_side(i,1) = cells_right(1,i); 
end
% if neither left side nor right side is complete, need to keep doing this with a different cell 
count_time = 0; %count how many times it's in the while loop, can break out after cell_num/2 times 
while (any(right_side) && any(left_side))
    count_time = count_time + 1;
    if count_time >= cell_num/2
        break %break out of while loop
    end
    % now try the right side
    if right_side(count_time,1) == 0 %in case there is no cell, - break immediately 
        break
    end
    TF_right_plus = SixteenCell_f(:,right_side(count_time,1)+2) == 1;
    TF_right_minus = SixteenCell_r(:,right_side(count_time,1)+2) == 1;
    TF_right = TF_right_plus | TF_right_minus;
    % if any other cell has any of these bins -> on the left
    cells_on_left_bin = SixteenCell_f(TF_right,3:end) | SixteenCell_r(TF_right,3:end);
    if size(cells_on_left_bin,1) == 1
        cells_on_left = cells_on_left_bin;
    else
        cells_on_left = any(cells_on_left_bin);
    end
    % remove the original cell
    cells_on_left(1,right_side(count_time,1)) = 0;
    cells_left = find(cells_on_left == 1);
    % find unique cells
    cells_left_unique = setdiff(cells_left,left_side);
    % find the first zero row
    non_zero_row = find(left_side==0,1,'first');
    for i = 1:size(cells_left_unique,2)
        left_side(non_zero_row+i-1,1) = cells_left_unique(i);
    end
    if  all(left_side) ~= 0 %done
        break
    else %then try the left side next cell
        if left_side(count_time+1,1) == 0
            break
        end  
        TF_left_plus = SixteenCell_f(:,left_side(count_time+1,1)+2) == 1;
        TF_left_minus = SixteenCell_r(:,left_side(count_time+1,1)+2) == 1;
        TF_left = TF_left_plus | TF_left_minus;
        % if any other cell has any of these bins -> on the right
        cells_on_right_bin = SixteenCell_f(TF_left,3:end) | SixteenCell_r(TF_left,3:end);
        cells_on_right = any(cells_on_right_bin);
        % remove the original cell
        cells_on_right(1,left_side(count_time+1,1)) = 0;
        cells_right = find(cells_on_right == 1);
        % find unique cells
        cells_right_unique = setdiff(cells_right,right_side);
        % find the first zero row
        non_zero_row = find(right_side==0,1,'first');
        for i = 1:size(cells_right_unique,2)
            right_side(non_zero_row+i-1,1) = cells_right_unique(i);
        end
    end
end
% if either side has all slots fill, can have the other side too
if all(left_side) ~= 0 || all(right_side) ~= 0
    if all(left_side) ~= 0
        left_side = left_side';
        right_side = setdiff(v,left_side); 
    else
        right_side = right_side';
        left_side = setdiff(v,right_side); 
    end
else %need to make different sets of left side and right side
    left_side = left_side';
    right_side = right_side';
    % find the cells leftover 
    cell_over_holder = setdiff(v,left_side); 
    cell_over = setdiff(cell_over_holder,right_side);
    % then need to distribute cell_over to the rest 
    % fill left side - then do setdiff to get right side
    left_side_nz = left_side; %non zero value
    left_side_nz(left_side_nz==0) = [];
    % see how many cases 
    over_cells = nchoosek(cell_over,cell_num/2-size(left_side_nz,2)); %choose cells for the left side 
    over_case = size(over_cells,1); 
    left_side_c = zeros(over_case,cell_num/2); 
    right_side_c = zeros(over_case,cell_num/2);
    for j = 1:over_case
        left_side_c(j,1:size(left_side_nz,2)) = left_side_nz ; %left_side cases; get the non zero    
        left_side_c(j,size(left_side_nz,2)+1:size(left_side_nz,2)+size(over_cells,2)) = over_cells(j,:); % get the rest from the over_cells
        % then get the right side
        right_side_c(j,:) = setdiff(v,left_side_c(j,:));
    end
    %then make it the final left_side and right_side
    left_side = left_side_c;
    right_side = right_side_c; 
end
M1_count = 0;
Ptotal = zeros(size(left_side,1)*315*315,cell_num);
for i = 1:size(left_side,1) %side1_new and side2_new is the same size, can do both concurrently
    v1 = left_side(i,:);
    min_num = min(v1);
    M11 = nchoosek(v1,4);
    MF11 = M11(:,1)~=min_num;
    M11(MF11,:) = [];
    v2 = right_side(i,:);
    min_num2 = min(v2); %same min_num should be ok -- make them different just in case
    M21 = nchoosek(v2,4);
    MF21 = M21(:,1)~=min_num2;
    M21(MF21,:) = [];
    for j = 1:size(M11,1)
        %then for each row of M11, M12, M21, M22 get N sets
        M12 = setdiff(v1,M11(j,:)); %M12 is just one row
        min_num3 = min(M11(j,:));
        N111 = nchoosek(M11(j,:),2);
        NF111 = N111(:,1)~=min_num3;
        N111(NF111,:) = [];
        min_num4 = min(M12);
        N121 = nchoosek(M12,2);
        NF121 = N121(:,1)~=min_num4;
        N121(NF121,:) = [];
        for k = 1:size(M21,1) %should be the same size of 35
            M22 = setdiff(v2,M21(k,:));
            min_num5 = min(M21(k,:));
            N211 = nchoosek(M21(k,:),2);
            NF211 = N211(:,1)~=min_num5;
            N211(NF211,:) = [];
            min_num6 = min(M22);
            N221 = nchoosek(M22,2);
            NF221 = N221(:,1)~=min_num6;
            N221(NF221,:) = [];
            for l = 1:size(N111) %should four should be 3
                N112 = setdiff(M11(j,:),N111(l,:));
                for m = 1:size(N121)
                    N122 = setdiff(M12,N121(m,:));
                    for n = 1:size(N211)
                        N212 = setdiff(M21(k,:),N211(n,:));
                        for o = 1:size(N221)
                            N222 = setdiff(M22,N221(o,:));
                            M1_count = M1_count + 1;
                            Ptotal(M1_count,1:2) = N111(l,:);
                            Ptotal(M1_count,3:4) = N112;
                            Ptotal(M1_count,5:6) = N121(m,:);
                            Ptotal(M1_count,7:8) = N122;
                            Ptotal(M1_count,9:10) = N211(n,:);
                            Ptotal(M1_count,11:12) = N212;
                            Ptotal(M1_count,13:14) = N221(o,:);
                            Ptotal(M1_count,15:16) = N222;
                        end
                    end
                end
            end
        end
    end
end
```
