## Supplementary material for "A probabilistic framework for cellular lineage reconstruction using single-cell 5-hydroxymethylcytosine sequencing": Code (scPECLR): Prediction_8.m

```
function [tree_result, fin_result] = Prediction_8(Cell_f, Cell_r, Ptotal, totbin, min_b, iter_max, tree_max)
% iterative prediction of 8 cell
[brk_num, b_obs] = Determine_Break_Num(Cell_f, Cell_r); %determine the break num
b_obs = max(b_obs,min_b); %set a minimum
b_3cell = [b_obs; b_obs; b_obs;];
expP_sort = Reconstruct_8_Tree(b_3cell, Ptotal, Cell_f, Cell_r);
% find the tree that has the higehst prob
iter_num = 1;
tree_result = zeros(iter_num,2*tree_max);
ii_max = zeros(tree_max,1);
for i = 1:tree_max
    ii_max(i) = expP_sort(i,2);
end
for j = 1:tree_max
    tree_result(iter_num,2*(j-1)+1) = ii_max(j);
    tree_result(iter_num,2*j) = expP_sort(j,1);
end
fin_result(iter_num,1) = ii_max(1); % the max tree
fin_result(iter_num,2) = b_obs;
fin_result(iter_num,3) = b_obs;
fin_result(iter_num,4) = b_obs;
% with the predicted tree; trace back to 4-cell and 2-cell
predict_tree = Ptotal(ii_max(1),:);
FourCell_re_f = zeros(sum(totbin),6); %4-cell reconstructed
FourCell_re_r = zeros(sum(totbin),6); %4-cell reconstructed
TwoCell_re_f = zeros(sum(totbin),4); %4-cell reconstructed
TwoCell_re_r = zeros(sum(totbin),4); %4-cell reconstructed
FourCell_re_f(:,1:2) = Cell_f(:,1:2);
FourCell_re_r(:,1:2) = Cell_f(:,1:2);
TwoCell_re_f(:,1:2) = Cell_f(:,1:2);
TwoCell_re_r(:,1:2) = Cell_f(:,1:2);
% make 4 cell
for i = 1:4
    % get the tree according to predicted topology
    first_cell = predict_tree((i-1)*2+1);
    second_cell = predict_tree(i*2);
    FourCell_re_f(:,2+i) = Cell_f(:,2+first_cell) + Cell_f(:,2+second_cell);
    FourCell_re_r(:,2+i) = Cell_r(:,2+first_cell) + Cell_r(:,2+second_cell);
end
for j = 1:2
    % this is ok since reconstructing from the reconstructed tree
    % already
    TwoCell_re_f(:,2+j) = FourCell_re_f(:,2*j+1) + FourCell_re_f(:,2*j+2);
    TwoCell_re_r(:,2+j) = FourCell_re_r(:,2*j+1) + FourCell_re_r(:,2*j+2);
end
% now calculate the break number in each cell division
[brk_num2, ~] = Determine_Break_Num(TwoCell_re_f,TwoCell_re_r); %get #break at 2-cell stage
[brk_num4, ~] = Determine_Break_Num(FourCell_re_f,FourCell_re_r); %get #break at 4-cell stage
% break at 8-cell stage from brk_num
brk_num4_8 = brk_num - brk_num4;
brk_num2_4 = brk_num4 - brk_num2;
brk_num1_2 = brk_num2;
b_re4_8 = poissfit([brk_num4_8(:,2); brk_num4_8(:,3)]);
b_re2_4 = poissfit([brk_num2_4(:,2); brk_num2_4(:,3)]);
b_re1_2 = poissfit([brk_num1_2(:,2); brk_num1_2(:,3)]);
b_re4_8 = max(b_re4_8,min_b);
b_re2_4 = max(b_re2_4,min_b);
b_re1_2 = max(b_re1_2,min_b);
% make a new b_3cell
b_3cell_new = [b_re1_2; b_re2_4; b_re4_8;];
% repredict the tree
expP_sort_new = Reconstruct_8_Tree(b_3cell_new, Ptotal, Cell_f,Cell_r);
% find the tree that has the higehst prob
iter_num = 2;
ii_max_new = zeros(tree_max,1); 
for i = 1:tree_max
    ii_max_new(i) = expP_sort_new(i,2);
end
for j = 1:tree_max
    tree_result(iter_num,2*(j-1)+1) = ii_max_new(j);
    tree_result(iter_num,2*j) = expP_sort_new(j,1);
end
fin_result(iter_num,1) = ii_max_new(1);  % the max tree
fin_result(iter_num,2) = b_re1_2;
fin_result(iter_num,3) = b_re2_4;
fin_result(iter_num,4) = b_re4_8;
while ii_max_new(1) ~= ii_max(1) && iter_num <= iter_max
    ii_max = ii_max_new;
    iter_num = iter_num + 1;
    % recalculate the whole thing
    ii_1max_new = expP_sort_new(1,2);
    % with the predicted tree; trace back to 4-cell and 2-cell
    predict_tree = Ptotal(ii_1max_new,:);
    FourCell_re_f = zeros(sum(totbin),6); %4-cell reconstructed
    FourCell_re_r = zeros(sum(totbin),6); %4-cell reconstructed
    TwoCell_re_f = zeros(sum(totbin),4); %2-cell reconstructed
    TwoCell_re_r = zeros(sum(totbin),4); %2-cell reconstructed
    FourCell_re_f(:,1:2) = Cell_f(:,1:2);
    FourCell_re_r(:,1:2) = Cell_f(:,1:2);
    TwoCell_re_f(:,1:2) = Cell_f(:,1:2);
    TwoCell_re_r(:,1:2) = Cell_f(:,1:2);
    % make 4 cell
    for i = 1:4
        % get the tree according to predicted topology
        first_cell = predict_tree((i-1)*2+1);
        second_cell = predict_tree(i*2);
        FourCell_re_f(:,2+i) = Cell_f(:,2+first_cell) + Cell_f(:,2+second_cell);
        FourCell_re_r(:,2+i) = Cell_r(:,2+first_cell) + Cell_r(:,2+second_cell);
    end
    for j = 1:2
        % this is ok since reconstructing from the reconstructed tree
        % already
        TwoCell_re_f(:,2+j) = FourCell_re_f(:,2*j+1) + FourCell_re_f(:,2*j+2);
        TwoCell_re_r(:,2+j) = FourCell_re_r(:,2*j+1) + FourCell_re_r(:,2*j+2);
    end
    % now calculate the break number in each cell division
    [brk_num2_new, ~] = Determine_Break_Num(TwoCell_re_f,TwoCell_re_r); %get #break at 2-cell stage
    [brk_num4_new, ~] = Determine_Break_Num(FourCell_re_f,FourCell_re_r); %get #break at 4-cell stage
    % break at 8-cell stage from brk_num
    brk_num4_8_new = brk_num - brk_num4_new;
    brk_num2_4_new = brk_num4_new - brk_num2_new;
    brk_num1_2_new = brk_num2_new;
    b_re4_8_new = poissfit([brk_num4_8_new(:,2); brk_num4_8_new(:,3)]);
    b_re2_4_new = poissfit([brk_num2_4_new(:,2); brk_num2_4_new(:,3)]);
    b_re1_2_new = poissfit([brk_num1_2_new(:,2); brk_num1_2_new(:,3)]);
    b_re4_8_new = max(b_re4_8_new,min_b);
    b_re2_4_new = max(b_re2_4_new,min_b);
    b_re1_2_new = max(b_re1_2_new,min_b);
    % make a new b_3cell
    b_3cell_new = [b_re1_2_new; b_re2_4_new; b_re4_8_new;];
    % repredict the tree
    expP_sort_new = Reconstruct_8_Tree(b_3cell_new, Ptotal, EightCell_f,Cell_r);
    ii_1max_new = expP_sort_new(1,2);
    ii_2max_new = expP_sort_new(2,2);
    ii_3max_new = expP_sort_new(3,2);
    tree_result(iter_num,1) = ii_1max_new;
    tree_result(iter_num,2) = expP_sort_new(1,1);
    tree_result(iter_num,3) = ii_2max_new;
    tree_result(iter_num,4) = expP_sort_new(2,1);
    tree_result(iter_num,5) = ii_3max_new;
    tree_result(iter_num,6) = expP_sort_new(3,1);
    fin_result(iter_num,1) = ii_1max_new; % the max tree
    fin_result(iter_num,2) = b_re1_2_new;
    fin_result(iter_num,3) = b_re2_4_new;
    fin_result(iter_num,4) = b_re4_8_new;
end
end
```
