## Supplementary material for "A probabilistic framework for cellular lineage reconstruction using single-cell 5-hydroxymethylcytosine sequencing": Code (scPECLR): Prediction_16.m

```
function [tree_result,fin_result] = Prediction_16(Cell_f ,Cell_r, Ptotal, totbin, min_b, iter_max, tree_max)
% iterative prediction of 16 cell
[~, b_obs] = Determine_Break_Num(Cell_f, Cell_r);
b_obs = max(b_obs,min_b);
b_4cell = [b_obs; b_obs; b_obs; b_obs;];
[expP_sort] = Reconstruct_16_Tree(b_4cell, Ptotal, Cell_f, Cell_r);
% find the tree that has the higehst prob
ii_max = zeros(tree_max,1);
for i = 1:tree_max
    ii_max(i) = expP_sort(i,2);
end
iter_num = 1;
tree_result = zeros(iter_num,2*tree_max);
for j = 1:tree_max
    tree_result(iter_num,2*(j-1)+1) = ii_max(j);
    tree_result(iter_num,2*j) = expP_sort(j,1);
end
fin_result(iter_num,1) = ii_max(1); % the max tree
fin_result(iter_num,2) = b_obs;
fin_result(iter_num,3) = b_obs;
fin_result(iter_num,4) = b_obs;
fin_result(iter_num,5) = b_obs;
% with the predicted tree; trace back to 8-cell, 4-cell and 2-cell
predict_tree = Ptotal(ii_max(1),:);
EightCell_re_f = zeros(sum(totbin),10); %8-cell reconstruct
EightCell_re_r = zeros(sum(totbin),10);
FourCell_re_f = zeros(sum(totbin),6); %4-cell reconstructed
FourCell_re_r = zeros(sum(totbin),6);
TwoCell_re_f = zeros(sum(totbin),4);
TwoCell_re_r = zeros(sum(totbin),4);
EightCell_re_f(:,1:2) = Cell_f(:,1:2);
EightCell_re_r(:,1:2) = Cell_f(:,1:2);
FourCell_re_f(:,1:2) = Cell_f(:,1:2);
FourCell_re_r(:,1:2) = Cell_f(:,1:2);
TwoCell_re_f(:,1:2) = Cell_f(:,1:2);
TwoCell_re_r(:,1:2) = Cell_f(:,1:2);
% make 8 cell
for i = 1:8
    % get the tree according to predicted topology
    first_cell = predict_tree((i-1)*2+1);
    second_cell = predict_tree(i*2);
    EightCell_re_f(:,2+i) = Cell_f(:,2+first_cell) + Cell_f(:,2+second_cell);
    EightCell_re_r(:,2+i) = Cell_r(:,2+first_cell) + Cell_r(:,2+second_cell);
end
for j = 1:4
    % this is ok since reconstructing from the reconstructed tree already
    FourCell_re_f(:,2+j) = EightCell_re_f(:,2*j+1) + EightCell_re_f(:,2*j+2);
    FourCell_re_r(:,2+j) = EightCell_re_r(:,2*j+1) + EightCell_re_r(:,2*j+2);
end
for k = 1:2
    TwoCell_re_f(:,2+k) = FourCell_re_f(:,2*k+1) + FourCell_re_f(:,2*k+2);
    TwoCell_re_r(:,2+k) = FourCell_re_r(:,2*k+1) + FourCell_re_r(:,2*k+2);
end
% now calculate the break number in each cell division
[brk_num2, ~] = Determine_Break_Num(TwoCell_re_f,TwoCell_re_r); %get #break at 2-cell stage
[brk_num4, ~] = Determine_Break_Num(FourCell_re_f,FourCell_re_r); %get #break at 4-cell stage
[brk_num8, ~] = Determine_Break_Num(EightCell_re_f,EightCell_re_r); %get #break at 8-cell stage
[brk_num16, ~] = Determine_Break_Num(Cell_f, Cell_r); %get #break at 16-cell stage
% find the break at each cell division
brk_num8_16 = brk_num16 - brk_num8;
brk_num4_8 = brk_num8 - brk_num4;
brk_num2_4 = brk_num4 - brk_num2;
brk_num1_2 = brk_num2;
b_re8_16 = poissfit([brk_num8_16(:,2); brk_num8_16(:,3)]);
b_re4_8 = poissfit([brk_num4_8(:,2); brk_num4_8(:,3)]);
b_re2_4 = poissfit([brk_num2_4(:,2); brk_num2_4(:,3)]);
b_re1_2 = poissfit([brk_num1_2(:,2); brk_num1_2(:,3)]);
b_re8_16 = max(b_re8_16,min_b);
b_re4_8 = max(b_re4_8,min_b);
b_re2_4 = max(b_re2_4,min_b);
b_re1_2 = max(b_re1_2,min_b);
% make a new b_3cell
b_4cell_new = [b_re1_2; b_re2_4; b_re4_8; b_re8_16;];
% repredict the tree
[expP_sort_new] = Reconstruct_16_Tree(b_4cell_new, Ptotal, Cell_f, Cell_r);
% find the tree that has the higehst prob
iter_num = 2;
ii_max_new = zeros(tree_max,1);
for i = 1:tree_max
    ii_max_new(i) = expP_sort_new(i,2);
end
for j = 1:tree_max
    tree_result(iter_num,2*(j-1)+1) = ii_max_new(j);
    tree_result(iter_num,2*j) = expP_sort_new(j,1);
end
fin_result(iter_num,1) = ii_max_new(1); % the max tree
fin_result(iter_num,2) = b_re1_2;
fin_result(iter_num,3) = b_re2_4;
fin_result(iter_num,4) = b_re4_8;
fin_result(iter_num,5) = b_re8_16;
while ii_max_new(1) ~= ii_max(1) && iter_num <= iter_max
    ii_max = ii_max_new;
    iter_num = iter_num + 1;
    % recalculate the whole thing
    ii_1max_new = expP_sort_new(1,2); %the predict tree
    % with the predicted tree; trace back to 8-cell, 4-cell and 2-cell
    predict_tree = Ptotal(ii_1max_new,:);
    EightCell_re_f = zeros(sum(totbin),10); %8-cell reconstructed
    EightCell_re_r = zeros(sum(totbin),10);
    FourCell_re_f = zeros(sum(totbin),6); %4-cell reconstructed
    FourCell_re_r = zeros(sum(totbin),6);
    TwoCell_re_f = zeros(sum(totbin),4);
    TwoCell_re_r = zeros(sum(totbin),4);
    EightCell_re_f(:,1:2) = Cell_f(:,1:2);
    EightCell_re_r(:,1:2) = Cell_f(:,1:2);
    FourCell_re_f(:,1:2) = Cell_f(:,1:2);
    FourCell_re_r(:,1:2) = Cell_f(:,1:2);
    TwoCell_re_f(:,1:2) = Cell_f(:,1:2);
    TwoCell_re_r(:,1:2) = Cell_f(:,1:2);
    for i = 1:8
        first_cell = predict_tree((i-1)*2+1);
        second_cell = predict_tree(i*2);
        EightCell_re_f(:,2+i) = Cell_f(:,2+first_cell) + Cell_f(:,2+second_cell);
        EightCell_re_r(:,2+i) = Cell_r(:,2+first_cell) + Cell_r(:,2+second_cell);
    end
    for j = 1:4
        FourCell_re_f(:,2+j) = EightCell_re_f(:,2*j+1) + EightCell_re_f(:,2*j+2);
        FourCell_re_r(:,2+j) = EightCell_re_r(:,2*j+1) + EightCell_re_r(:,2*j+2);
    end
    for k = 1:2
        TwoCell_re_f(:,2+k) = FourCell_re_f(:,2*k+1) + FourCell_re_f(:,2*k+2);
        TwoCell_re_r(:,2+k) = FourCell_re_r(:,2*k+1) + FourCell_re_r(:,2*k+2);
    end
    [brk_num2_new, ~] = Determine_Break_Num(TwoCell_re_f,TwoCell_re_r); %get #break at 2-cell stage
    [brk_num4_new, ~] = Determine_Break_Num(FourCell_re_f,FourCell_re_r); %get #break at 4-cell stage
    [brk_num8_new, ~] = Determine_Break_Num(EightCell_re_f,EightCell_re_r); %get #break at 4-cell stage
    brk_num8_16_new = brk_num16 - brk_num8_new;
    brk_num4_8_new = brk_num8_new - brk_num4_new;
    brk_num2_4_new = brk_num4_new - brk_num2_new;
    brk_num1_2_new = brk_num2_new;
    b_re8_16_new = poissfit([brk_num8_16_new(:,2); brk_num8_16_new(:,3)]);
    b_re4_8_new = poissfit([brk_num4_8_new(:,2); brk_num4_8_new(:,3)]);
    b_re2_4_new = poissfit([brk_num2_4_new(:,2); brk_num2_4_new(:,3)]);
    b_re1_2_new = poissfit([brk_num1_2_new(:,2); brk_num1_2_new(:,3)]);
    b_re8_16_new = max(b_re8_16_new,min_b);
    b_re4_8_new = max(b_re4_8_new,min_b);
    b_re2_4_new = max(b_re2_4_new,min_b);
    b_re1_2_new = max(b_re1_2_new,min_b);
    b_4cell_new = [b_re1_2_new; b_re2_4_new; b_re4_8_new; b_re8_16_new;];
    % repredict the tree
    [expP_sort_new] = Reconstruct_16_Tree(b_4cell_new, Ptotal, Cell_f, Cell_r);
    for i = 1:tree_max
        ii_max_new(i) = expP_sort_new(i,2);
    end
    for j = 1:tree_max
        tree_result(iter_num,2*(j-1)+1) = ii_max_new(j);
        tree_result(iter_num,2*j) = expP_sort_new(j,1);
    end
    fin_result(iter_num,1) = ii_max_new(1); % the max tree
    fin_result(iter_num,2) = b_re1_2_new;
    fin_result(iter_num,3) = b_re2_4_new;
    fin_result(iter_num,4) = b_re4_8_new;
    fin_result(iter_num,4) = b_re8_16_new;
end
end
```
