## Supplementary material for "A probabilistic framework for cellular lineage reconstruction using single-cell 5-hydroxymethylcytosine sequencing": Code (scPECLR): Prob_Cal.m

```
function [com_f, com_r, logP] = Prob_Cal(f1, f2, r1, r2, b)
% calculate probability (in -ln) of 2 cells 5hmC from the same mother cell 
% input: forward and reverse strands of both cells (for a chromosome of interest) and SCE rate  
% output: combined forward and reverse strand and -ln of the probability
totbin = size(f1,1);
N = totbin - 1;
r = 0.5;
% create the forward and reverse strands of the mother cell 
com_f = f1 + f2;
com_r = r1 + r2;
com_t = com_f + com_r;
if sum(com_t) == 0 %% if there is nothing, then skip that iteration
    prob = 1;
    logP = -log(prob);
    return % skip the rest of the code
end
prob = r/exp(b); % start off with r/exp(b)
% if the whole strand (plus or minus) does not break at all and passes on then r/e^b and skips that iteration
% do not consider even breaks occur at the exact same locations 
if (sum(com_f) == totbin && (sum(f1) == totbin || sum(f2) == totbin)) || ...
        (sum(com_r) == totbin && (sum(r1) == totbin || sum(r2) == totbin))
    logP = -log(prob);
    return
end
% most of the times, there would only be + or - strand so we break down to
% 3 cases: + only, - only and + or -
if sum(com_r) == 0 %nothing in the minus strand, essentially just +
    % break in the combined cell
    clear break_count; % 1 = cell1, 2 = cell2
    num_com = 0;
    num_1 = 0;
    num_2 = 0;
    af_com = com_f(1);
    af_1 = f1(1);
    af_2 = f2(1);
    piece_count = 1;
    piece(1,1) = af_com;
    piece(1,2) = 1;
    for k = 1:totbin % bins
        bf_com = com_f(k);
        if bf_com ~= af_com
            num_com = num_com+1;
            af_com = bf_com;
            piece_count = piece_count+1;    %also need to keep track of combine1's 5hmC distribution
            piece(piece_count,1) = af_com;
            piece(piece_count,2) = k; % where that piece starts
        end
        % count number of breaks in cell1 forward (but not necessarily with cell2) 
        bf_1 = f1(k);
        if bf_1 ~= af_1
            num_1 = num_1+1;
            af_1 = bf_1;
        end
        % count number of breaks in cell2 forward (but not necessarily with cell1) 
        bf_2 = f2(k);
        if bf_2 ~= af_2
            num_2 = num_2+1;
            af_2 = bf_2;
        end
    end
    break_count = (num_1 + num_2 - num_com)/2; %calculate break count that is shared between the 2 cells 
    if break_count > 0
        prob = prob*b^break_count/(N^break_count); % add the break term
    end
    % if the combine piece has black region at the end; add exponential term
    if piece(1,1) == 0 %left end
        R1 = (piece(2,2) - piece(1,2))/totbin;
    else
        R1 = 0;
    end
    if piece(size(piece,1),1) == 0 %right end
        Rnp1 = (totbin - piece(size(piece,1),2)+1)/totbin;
    else
        Rnp1 = 0;
    end
    Redge = R1 + Rnp1;
    if Redge > 0
        prob = prob * exp(b*Redge*(N+1)/N); %add exponetial term of the edge 
    end
    % black region in the middle; find all the black pieces in the middle
    mid_piece = [];
    for i = 2:size(piece,1)-1 %ignore first and last one
        if piece(i,1) == 0
            mid_piece = [mid_piece; i]; % add that row into the matrix
        end
    end
    % add the size of the piece
    if ~isempty(mid_piece)
        for j = 1:size(mid_piece,1)
            mp_row = mid_piece(j,1);
            mid_piece(j,2) = (piece(mp_row+1,2)-piece(mp_row,2))/totbin; % the length ratio of the middle piece 
            % if the 2 pieces beside the midpiece is in the same cell then it's a cosh
            % if they are different it's a sinh; to check that we need to only check the
            % last bin of the prior piece (where the chromosome starts + where the mid piece starts within that chr - 2)
            % and the first bin of the following piece (where the chr starts + where the next piece starts -1)
            % -1 is to discount the starts of first chromosome
            if (f1(1+piece(mp_row,2)-2,1) == 1 && f1(1+piece(mp_row+1,2)-1,1) == 1) || ...
                    (f2(1+piece(mp_row,2)-2,1) == 1 && f2(1+piece(mp_row+1,2)-1,1) == 1) %then this is cosh
                prob = prob * cosh((mid_piece(j,2)*(N+1)+1)*b/N); % add the term
            elseif (f1(1+piece(mp_row,2)-2,1) == 1 && f2(1+piece(mp_row+1,2)-1,1) == 1) || ...
                    (f2(1+piece(mp_row,2)-2,1) == 1 && f1(1+piece(mp_row+1,2)-1,1) == 1) %then this is sinh
                prob = prob * sinh((mid_piece(j,2)*(N+1)+1)*b/N);
            else
                msg = 'Error of determining the middle SCE occurred in the plus strand.';
                error(msg)
            end
        end
    end
elseif sum(com_f) == 0 %nothing in the plus strand, essentially just -
    clear break_count; % 1 = cell1, 2 = cell2
    num_com = 0;
    num_1 = 0;
    num_2 = 0;
    ar_com = com_r(1);
    ar_1 = r1(1);
    ar_2 = r2(1);
    piece_count = 1;
    piece(1,1) = ar_com;
    piece(1,2) = 1;
    for k = 1:totbin %bins
        br_com = com_r(k);
        if br_com ~= ar_com
            num_com = num_com+1;
            ar_com = br_com;
            piece_count = piece_count + 1; %need to keep track of combine1's 5hmC distribution
            piece(piece_count,1) = ar_com;
            piece(piece_count,2) = k; % where that piece starts
        end
        br_1 = r1(k);
        if br_1 ~= ar_1
            num_1 = num_1+1;
            ar_1 = br_1;
        end
        br_2 = r2(k);
        if br_2 ~= ar_2
            num_2 = num_2+1;
            ar_2 = br_2;
        end
    end
    break_count = (num_1 + num_2 - num_com)/2;
    if break_count > 0
        prob = prob*b^break_count/(N^break_count);
    end
    if piece(1,1) == 0
        R1 = (piece(2,2) - piece(1,2))/totbin;
    else
        R1 = 0;
    end
    if piece(size(piece,1),1) == 0
        Rnp1 = (totbin - piece(size(piece,1),2)+1)/totbin;
    else
        Rnp1 = 0;
    end
    Redge = R1 + Rnp1;
    if Redge > 0
        prob = prob * exp(b*Redge*(N+1)/N);
    end
    mid_piece = [];
    for i = 2:size(piece,1)-1 %ignore first and last one
        if piece(i,1) == 0
            mid_piece = [mid_piece; i]; % add that row into the matrix
        end
    end
    if ~isempty(mid_piece)
        for j = 1:size(mid_piece,1)
            mp_row = mid_piece(j,1);
            mid_piece(j,2) = (piece(mp_row+1,2)-piece(mp_row,2))/totbin;
            if (r1(1+piece(mp_row,2)-2,1) == 1 && r1(1+piece(mp_row+1,2)-1,1) == 1) || ...
                    (r2(1+piece(mp_row,2)-2,1) == 1 && r2(1+piece(mp_row+1,2)-1,1) == 1) %then this is cosh
                prob = prob * cosh((mid_piece(j,2)*(N+1)+1)*b/N);
            elseif (r1(1+piece(mp_row,2)-2,1) == 1 && r2(1+piece(mp_row+1,2)-1,1) == 1) || ...
                    (r2(1+piece(mp_row,2)-2,1) == 1 && r1(1+piece(mp_row+1,2)-1,1) == 1) %then this is sinh
                prob = prob * sinh((mid_piece(j,2)*(N+1)+1)*b/N);
            else
                msg = 'Error of determining the middle SCE occurred in the minus strand.';
                error(msg)
            end
        end
    end
else %then it has both + and - strands. 
    % there are 2 cases, full length on both strands (1 to 2 cells) but  
    % and the other cases (+/- strands cannot have the same bin)
    if all(com_t(:) == 2 ) == 1 % full strand on both sides - the mother is the 1-cell zygote
        %in this case the number of break at the exact spot will be the
        %same as the number of break found in one of the strand -- use f1
        clear break_count; % 1 = cell1, 2 = cell2
        num_1 = 0;
        af_1 = f1(1);
        for k = 1:totbin %bins
            bf_1 = f1(k);
            if bf_1 ~= af_1
                num_1 = num_1+1;
                af_1 = bf_1;
            end
        end
        break_count = num_1;
        if break_count > 0 %it should always be bigger than 0, because otherwise it would end a lot earlier
            prob = prob*b^break_count/(N^break_count);
        end
        logP = -log(prob);
        return;
    elseif any(com_t(:) == 2) == 1 %any of the bins have 5hmC in both +/- strand but both are not all FL
        msg = 'The 2 cells considered should not be in the same subtree.';
        error(msg)
    else %it has both + and - strands but neither is FL; then the number of exact break would be number in (4 daughter strands - number in 2 mother strands)/2
        clear break_count; % 1 = cell1, 2 = cell2
        num_com_f = 0;
        num_com_r = 0;
        num_com_t = 0;
        num_1_f = 0;
        num_1_r = 0;
        num_2_f = 0;
        num_2_r = 0;
        % need to keep track of every piece to track the special case
        af_com = com_f(1); %mother cell 
        pcfcom = 1;  
        pfcom(1,1) = af_com;
        pfcom(1,2) = 1;
        ar_com = com_r(1);
        pcrcom = 1; 
        prcom(1,1) = ar_com;
        prcom(1,2) = 1;
        af_1 = f1(1); %cell1 
        pcf1 = 1; 
        pf1(1,1) = af_1;
        pf1(1,2) = 1;
        ar_1 = r1(1);
        pcr1 = 1; 
        pr1(1,1) = ar_1;
        pr1(1,2) = 1;
        af_2 = f2(1); %cell2
        pcf2 = 1; 
        pf2(1,1) = af_2; 
        pf2(1,2) = 1;
        ar_2 = r2(1);
        pcr2 = 1; 
        pr2(1,1) = ar_2;
        pr2(1,2) = 1;
        at_com = com_t(1);
        piece_count = 1; %but the piece count will be from total combined count
        piece(1,1) = at_com;
        piece(1,2) = 1;
        for k = 1:totbin %bins
            % mother cell 
            bt_com = com_t(k);
            if bt_com ~= at_com
                num_com_t = num_com_t+1;
                at_com = bt_com;
                piece_count = piece_count+1;    % this piece will contain every single break location
                piece(piece_count,1) = at_com;
                piece(piece_count,2) = k; % where that piece starts
            end
            bf_com = com_f(k);
            if bf_com ~= af_com
                num_com_f = num_com_f+1;
                af_com = bf_com;
                pcfcom = pcfcom + 1;
                pfcom(pcfcom,1) = af_com;
                pfcom(pcfcom,2) = k;         
            end
            br_com = com_r(k);
            if br_com ~= ar_com
                num_com_r = num_com_r+1;
                ar_com = br_com;
                pcrcom = pcrcom + 1;
                prcom(pcrcom,1) = ar_com;
                prcom(pcrcom,2) = k;
            end
            % cell1
            bf_1 = f1(k);
            if bf_1 ~= af_1
                num_1_f = num_1_f+1;
                af_1 = bf_1;
                pcf1 = pcf1 + 1;
                pf1(pcf1,1) = af_1;
                pf1(pcf1,2) = k; 
            end
            br_1 = r1(k);
            if br_1 ~= ar_1
                num_1_r = num_1_r+1;
                ar_1 = br_1;
                pcr1 = pcr1 + 1;
                pr1(pcr1,1) = ar_1;
                pr1(pcr1,2) = k; 
            end
            % cell2
            bf_2 = f2(k);
            if bf_2 ~= af_2
                num_2_f = num_2_f+1;
                af_2 = bf_2;
                pcf2 = pcf2 + 1;
                pf2(pcf2,1) = af_2;
                pf2(pcf2,2) = k; 
            end
            br_2 = r2(k);
            if br_2 ~= ar_2
                num_2_r = num_2_r+1;
                ar_2 = br_2;
                pcr2 = pcr2 + 1;
                pr2(pcr2,1) = ar_2;
                pr2(pcr2,2) = k; 
            end
        end
        break_count = (num_1_f + num_1_r + num_2_f + num_2_r - num_com_f - num_com_r)/2;
        if break_count > 0
            prob = prob*b^break_count/(N^break_count);
        end
        % need a special case where the break at the exact spot keeping 
        % "continuous +/- strand" from mother cell to just one daughter cell - that counts as a break 
        % find in pf_com going from 1 to 0 or 0 to 1 and that part is
        % immediately pick up by the pr_com 0 to 1 or 1 to 0 respectively
        % since f_com and r_com cannot be 1 at the same time except at the 1 cell state
        % this can be done by checking if the start of the piece in pfcom and prcom overlaps 
        overlap_com = intersect(pfcom(:,2),prcom(:,2));
        overlap_com(overlap_com(:)==1,:) = []; % need to remove 1 out 
        if ~isempty(overlap_com)
            for i = 1:size(overlap_com,1)
                ov_st = overlap_com(i); % start position of the overlap
                if (any(pf1(:,2) == ov_st) && any(pr1(:,2) == ov_st)) || ...
                        (any(pf2(:,2) == ov_st) && any(pr2(:,2) == ov_st)) %either of the cell has the continuous +/- strands
                    prob = prob*b/N;
                end % ok if it's 2 different cells 
            end
        end     
        % infinite breaks can happen at the edge of the combined +/- strands
        if piece(1,1) == 0
            R1 = (piece(2,2) - piece(1,2))/totbin;
        else
            R1 = 0;
        end
        if piece(size(piece,1),1) == 0
            Rnp1 = (totbin - piece(size(piece,1),2)+1)/totbin;
        else
            Rnp1 = 0;
        end
        Redge = R1 + Rnp1;
        if Redge > 0
            prob = prob * exp(b*Redge*(N+1)/N);
        end
        mid_piece = [];
        for i = 2:size(piece,1)-1 %ignore first and last one
            if piece(i,1) == 0
                mid_piece = [mid_piece; i]; % add that row into the matrix
            end
        end
        if ~isempty(mid_piece)
            for j = 1:size(mid_piece,1)
                mp_row = mid_piece(j,1);
                mid_piece(j,2) = (piece(mp_row+1,2)-piece(mp_row,2))/totbin;
                % there are 4 cases;
                %   combine       same/different cell   term
                % 1 same strand   same cell             cosh
                % 2 same          different             sinh
                % 3 oppo          same                  sinh
                % 4 oppo          diff                  cosh
                % are the 2 pieces next to it on the same strand or opposite strand
                if (com_f(1+piece(mp_row,2)-2,1) == 1 && com_f(1+piece(mp_row+1,2)-1,1) == 1) %on the same forward
                    if (f1(1+piece(mp_row,2)-2,1) == 1 && f1(1+piece(mp_row+1,2)-1,1) == 1) || ...
                            (f2(1+piece(mp_row,2)-2,1) == 1 && f2(1+piece(mp_row+1,2)-1,1) == 1) %then this is cosh
                        prob = prob * cosh((mid_piece(j,2)*(N+1)+1)*b/N);
                    elseif (f1(1+piece(mp_row,2)-2,1) == 1 && f2(1+piece(mp_row+1,2)-1,1) == 1) || ...
                            (f2(1+piece(mp_row,2)-2,1) == 1 && f1(1+piece(mp_row+1,2)-1,1) == 1) %then this is sinh
                        prob = prob * sinh((mid_piece(j,2)*(N+1)+1)*b/N);
                    else
                        msg = 'Error of determining the middle SCE occurred in the plus strand.';
                        error(msg)
                    end
                elseif(com_r(1+piece(mp_row,2)-2,1) == 1 && com_r(1+piece(mp_row+1,2)-1,1) == 1) %on the same reverse
                    if (r1(1+piece(mp_row,2)-2,1) == 1 && r1(1+piece(mp_row+1,2)-1,1) == 1) || ...
                            (r2(1+piece(mp_row,2)-2,1) == 1 && r2(1+piece(mp_row+1,2)-1,1) == 1) %then this is cosh
                        prob = prob * cosh((mid_piece(j,2)*(N+1)+1)*b/N);
                    elseif (r1(1+piece(mp_row,2)-2,1) == 1 && r2(1+piece(mp_row+1,2)-1,1) == 1) || ...
                            (r2(1+piece(mp_row,2)-2,1) == 1 && r1(1+piece(mp_row+1,2)-1,1) == 1) %then this is sinh
                        prob = prob * sinh((mid_piece(j,2)*(N+1)+1)*b/N);
                    else
                        msg = 'Error of determining the middle SCE occurred in the minus strand.';
                        error(msg)
                    end
                elseif (com_f(1+piece(mp_row,2)-2,1) == 1 && com_r(1+piece(mp_row+1,2)-1,1) == 1) %left on forward, right on reverse
                    if (f1(1+piece(mp_row,2)-2,1) == 1 && r1(1+piece(mp_row+1,2)-1,1) == 1) || ...
                            (f2(1+piece(mp_row,2)-2,1) == 1 && r2(1+piece(mp_row+1,2)-1,1) == 1) %the same cell, then this is sinh
                        prob = prob * sinh((mid_piece(j,2)*(N+1)+1)*b/N);
                    elseif (f1(1+piece(mp_row,2)-2,1) == 1 && r2(1+piece(mp_row+1,2)-1,1) == 1) || ...
                            (f2(1+piece(mp_row,2)-2,1) == 1 && r1(1+piece(mp_row+1,2)-1,1) == 1) %different cell, cosh
                        prob = prob * cosh((mid_piece(j,2)*(N+1)+1)*b/N);
                    else
                        msg = 'Error of determining the middle SCE occurred.';
                        error(msg)
                    end
                elseif (com_r(1+piece(mp_row,2)-2,1) == 1 && com_f(1+piece(mp_row+1,2)-1,1) == 1) %left on reverse, right on forward
                    if (r1(1+piece(mp_row,2)-2,1) == 1 && f1(1+piece(mp_row+1,2)-1,1) == 1) || ...
                            (r2(1+piece(mp_row,2)-2,1) == 1 && f2(1+piece(mp_row+1,2)-1,1) == 1) %the same cell, then this is sinh
                        prob = prob * sinh((mid_piece(j,2)*(N+1)+1)*b/N);
                    elseif (r1(1+piece(mp_row,2)-2,1) == 1 && f2(1+piece(mp_row+1,2)-1,1) == 1) || ...
                            (r2(1+piece(mp_row,2)-2,1) == 1 && f1(1+piece(mp_row+1,2)-1,1) == 1) %different cell, cosh
                        prob = prob * cosh((mid_piece(j,2)*(N+1)+1)*b/N);
                    else
                        msg = 'Error of determining the middle SCE occurred.';
                        error(msg)
                    end
                else
                    msg = 'Error occurred.';
                    error(msg)
                end
                
            end
        end
        
    end
end
logP = -log(prob);
end
```
