## Supplementary material for "A probabilistic framework for cellular lineage reconstruction using single-cell 5-hydroxymethylcytosine sequencing": Code (scPECLR): readme.txt

The files are supported by MATLAB (tested in ver. 2018a).
One main test file provided: sc_PECLR_test.m
2 test data provided: test8cell_embryo.mat, test16cell_embryo.mat
test8cell_embryo.mat is the dataset used to generate Fig. 1b, contains EightCell_f, EightCell_r
test16cell_embryo.mat was generated with constant SCE rate of 0.8, contains SixteenCell_f, SixteenCell_r
scPECLR is the main function that calls other functions provided in the folder
  Determine_Break_Num: determines SCE events and transitions among the cells
  OSS_8/OSS_16: performs OSS analysis
  Prediction_8/Prediction_16: iteratively calculates probability of each tree topology
  Prob_Cal: calculates the probability ratio based on SCE pattern
  Reconstruct_8_Tree/Reconstruct_16_Tree: calculates probability of each tree topology (1 iteration)
[fin_result, tree_result, Ptotal] = scPECLR(forward_5hmC,reverse_5hmC)
fin_result shows the predicted tree (tree with the highest probability)
  Each row shows the result after a certain number of iteration (row1 = after 1 iteration)
  1st column: the tree topology. That row of Ptotal is the predicted topology.
  2nd-4th   : b1-b3 used to calculate the probability (2nd-5th for b1-b4 in 16-cell prediction)
tree_result : collects the trees with the highest probability
  Each row shows the result after a certain number of iteration
  The odd columns are the topologies and the even columns are their associated predicted probabilities
  Top 3 trees and top 10 trees are kept for 8-cell and 16-cell prediction respectively
Ptotal shows the topologies considered. Each row is one topology. Column 1 is sister with column 2
  and their mother is sister with the mother cell of columns 3 and 4, and so on.
