## Supplementary material for "A probabilistic framework for cellular lineage reconstruction using single-cell 5-hydroxymethylcytosine sequencing": Code (scPECLR): Reconstruct_8_Tree.m

```
function [expP_sort] = Reconstruct_8_Tree(b_3cell, Ptotal, EightCell_f, EightCell_r)
%%% Reconstruct 8 cells -- comes out with tree prediction
%%% Input: SCE rate for each cell division, Trees being considered EightCell - forward and reverse, th
%%% Output: expP_sort 1st col = prob 
%%% SHOULD USE FLS_EX OR OSS_8 FIRST
chr_num = max(EightCell_f(:,1));
start(1,1) = 1;
for i = 1:chr_num-1
    stop(i,1) = max(find(EightCell_f(:,1)==i));
    start(i+1,1) = stop(i,1)+1;
end
stop(chr_num,1) = size(EightCell_f,1);
PtotalLeft = unique(Ptotal(:,1:4),'rows');
PtotalRight = unique(Ptotal(:,5:8),'rows');
% left side first; get the 4 cells and make a 1:1 map between cell number
% and the column it will be in FourCell_f 
FourCell_f(:,1:2) = EightCell_f(:,1:2);
FourCell_r(:,1:2) = EightCell_r(:,1:2);
OnetoOne_map = zeros(4,2);
for i = 1:4
    OnetoOne_map(i,1) = i;
    OnetoOne_map(i,2) = PtotalLeft(1,i);
    FourCell_f(:,2+i) = EightCell_f(:,2+PtotalLeft(1,i));
    FourCell_r(:,2+i) = EightCell_r(:,2+PtotalLeft(1,i));
end
Prem = PtotalLeft; % use a variable with a constant name Prem 
% create an original struct of 2-cell state
% all the pairs of cells combined to 2 cell state
T2l = struct('cell1', struct, 'cell2', struct);
for i = 1:size(Prem,1)
    for j = 1:chr_num
        % cells 1/2
        cell1 = Prem(i,1); %cell they want
        cell2 = Prem(i,2); 
        col1 = find(OnetoOne_map(:,2) == cell1); %column in the four cell they want
        col2 = find(OnetoOne_map(:,2) == cell2);
        f1 = FourCell_f(start(j):stop(j),col1+2); %start and stop of each chromosome; column of cell 1
        f2 = FourCell_f(start(j):stop(j),col2+2);
        r1 = FourCell_r(start(j):stop(j),col1+2);
        r2 = FourCell_r(start(j):stop(j),col2+2);
        [com_f, com_r, logP ] = Prob_Cal(f1, f2, r1, r2, b_3cell(3));
        T2l(i).col1(j).F = com_f;
        T2l(i).col1(j).R = com_r;
        T2l(i).col1(j).P = logP;
        % cells 3/4
        cell1 = Prem(i,3); %cell they want
        cell2 = Prem(i,4); 
        col1 = find(OnetoOne_map(:,2) == cell1); %column in the four cell they want
        col2 = find(OnetoOne_map(:,2) == cell2);
        f1 = FourCell_f(start(j):stop(j),col1+2); %start and stop of each chromosome; column of cell 1
        f2 = FourCell_f(start(j):stop(j),col2+2);
        r1 = FourCell_r(start(j):stop(j),col1+2);
        r2 = FourCell_r(start(j):stop(j),col2+2);
        [com_f, com_r, logP ] = Prob_Cal(f1, f2, r1, r2, b_3cell(3));
        T2l(i).col2(j).F = com_f;
        T2l(i).col2(j).R = com_r;
        T2l(i).col2(j).P = logP;
    end
end
% make to 1 cell stage
T1l = struct;
for i = 1:size(Prem,1)
    for j = 1:chr_num
        f1 = T2l(i).col1(j).F;
        f2 = T2l(i).col2(j).F;
        r1 = T2l(i).col1(j).R;
        r2 = T2l(i).col2(j).R;
        [com_f, com_r, logP ] = Prob_Cal( f1, f2, r1, r2, b_3cell(2));
        T1l(i).col1(j).F = com_f;
        T1l(i).col1(j).R = com_r;
        T1l(i).col1(j).P = logP;
    end
end
totPl = zeros(size(Prem,1),1);
% then for each tree sum all P
for i = 1:size(Prem,1) %save totPl for each case; then cross all the cases at once outside time for loop
    for j = 1:chr_num
        totPl(i,1) = totPl(i,1) + T2l(i).col1(j).P + T2l(i).col2(j).P + T1l(i).col1(j).P;
    end
end

% right side 
FourCell_right_f(:,1:2) = EightCell_f(:,1:2);
FourCell_right_r(:,1:2) = EightCell_r(:,1:2);
OnetoOne_map = zeros(4,2);
for i = 1:4
    OnetoOne_map(i,1) = i;
    OnetoOne_map(i,2) = PtotalRight(1,i);
    FourCell_right_f(:,2+i) = EightCell_f(:,2+PtotalRight(1,i));
    FourCell_right_r(:,2+i) = EightCell_r(:,2+PtotalRight(1,i));
end
Prem = PtotalRight;
T2r = struct('cell1', struct, 'cell2', struct);
for i = 1:size(Prem,1)
    for j = 1:chr_num
        cell1 = Prem(i,1); %cell they want
        cell2 = Prem(i,2); 
        col1 = find(OnetoOne_map(:,2) == cell1); %column in the four cell they want
        col2 = find(OnetoOne_map(:,2) == cell2);
        f1 = FourCell_right_f(start(j):stop(j),col1+2); %start and stop of each chromosome; column of cell 1
        f2 = FourCell_right_f(start(j):stop(j),col2+2);
        r1 = FourCell_right_r(start(j):stop(j),col1+2);
        r2 = FourCell_right_r(start(j):stop(j),col2+2);
        [com_f, com_r, logP ] = Prob_Cal( f1, f2, r1, r2, b_3cell(3));
        T2r(i).col1(j).F = com_f;
        T2r(i).col1(j).R = com_r;
        T2r(i).col1(j).P = logP;
        cell1 = Prem(i,3); %cell they want
        cell2 = Prem(i,4); 
        col1 = find(OnetoOne_map(:,2) == cell1); %column in the four cell they want
        col2 = find(OnetoOne_map(:,2) == cell2);
        f1 = FourCell_right_f(start(j):stop(j),col1+2); %start and stop of each chromosome; column of cell 1
        f2 = FourCell_right_f(start(j):stop(j),col2+2);
        r1 = FourCell_right_r(start(j):stop(j),col1+2);
        r2 = FourCell_right_r(start(j):stop(j),col2+2);
        [com_f, com_r, logP ] = Prob_Cal(f1, f2, r1, r2, b_3cell(3));
        T2r(i).col2(j).F = com_f;
        T2r(i).col2(j).R = com_r;
        T2r(i).col2(j).P = logP;
    end
end
T1r = struct; 
for i = 1:size(Prem,1)
    for j = 1:chr_num
        f1 = T2r(i).col1(j).F;
        f2 = T2r(i).col2(j).F;
        r1 = T2r(i).col1(j).R;
        r2 = T2r(i).col2(j).R;
        [com_f, com_r, logP ] = Prob_Cal(f1, f2, r1, r2, b_3cell(2));
        T1r(i).col1(j).F = com_f;
        T1r(i).col1(j).R = com_r;
        T1r(i).col1(j).P = logP;
    end
end
% then for each tree sum all P
totPr = zeros(size(Prem,1),1);
for i = 1:size(Prem,1)
    for j = 1:chr_num
        totPr(i,1) = totPr(i,1) + T2r(i).col1(j).P + T2r(i).col2(j).P + T1r(i).col1(j).P;
    end
end
row_totP = 1;
for o = 1:size(Prem,1)
    for p = 1:size(Prem,1)
        totP(row_totP,1) = totPl(o) + totPr(p);
        row_totP = row_totP + 1;       
    end
end
totP2 = -totP;
totP2 = totP2 - max(totP2)+100; %normalized the max one to 100
expP = exp(totP2);
expP_sum = sum(expP);
expP_prob = expP /expP_sum;
expP_prob(1:9,2) = 1:9;
expP_sort = sortrows(expP_prob,'descend');
end
```
