## Supplementary material for "A probabilistic framework for cellular lineage reconstruction using single-cell 5-hydroxymethylcytosine sequencing": Code (scPECLR): Reconstruct_16_Tree.m

```
function [expP_sort_top] = Reconstruct_16_Tree(b_4cell,  Ptotal, SixteenCell_f, SixteenCell_r)
%%% Reconstruct 16 cells -- comes out with tree prediction
%%% Input: SCE rate for each cell division SixteenCell, Ptotal, forward and reverse strands
%%% Output: expP_sort 1st col = prob, 2nd col = prob and Ptotal16
% NOTE THAT B RATE FOR THE FIRST CELL DIV IS NOT USED
% left side first -- essentially 8cell
max_num = 100; % take only top 100 trees
chr_num = max(SixteenCell_f(:,1));
start(1,1) = 1;
stop = zeros(chr_num,1);
for i = 1:chr_num-1
    stop(i,1) = find(SixteenCell_f(:,1)==i, 1, 'last');
    start(i+1,1) = stop(i,1)+1;
end
stop(chr_num,1) = size(SixteenCell_f,1);
PtotalLeft = unique(Ptotal(:,1:8),'rows');
PtotalRight = unique(Ptotal(:,9:16),'rows');
Premove = PtotalLeft;
% create an original struct of 4-cell state
% all the pairs of cells combined to 4 cell state
T4l = struct('cell1', struct, 'cell2', struct,'cell3', struct,'cell4', struct);
for i = 1:size(Premove,1)
    EightCell_left_f(:,1:10) = SixteenCell_f(:,[1:2 PtotalLeft(i,:)+2]); %make a new 8 cell left in each Ptotalleft case 
    EightCell_left_r(:,1:10) = SixteenCell_r(:,[1:2 PtotalLeft(i,:)+2]);
    for j = 1:chr_num
        f1 = EightCell_left_f(start(j):stop(j),3); %start and stop of each chromosome; column of cell 1
        f2 = EightCell_left_f(start(j):stop(j),4);
        r1 = EightCell_left_r(start(j):stop(j),3);
        r2 = EightCell_left_r(start(j):stop(j),4);
        [com_f, com_r, logP ] = Prob_Cal(f1, f2, r1, r2, b_4cell(4));
        T4l(i).cell1(j).F = com_f;
        T4l(i).cell1(j).R = com_r;
        T4l(i).cell1(j).P = logP;
        f1 = EightCell_left_f(start(j):stop(j),5); %start and stop of each chromosome; column of cell 1
        f2 = EightCell_left_f(start(j):stop(j),6);
        r1 = EightCell_left_r(start(j):stop(j),5);
        r2 = EightCell_left_r(start(j):stop(j),6);
        [com_f, com_r, logP ] = Prob_Cal(f1, f2, r1, r2, b_4cell(4));
        T4l(i).cell2(j).F = com_f;
        T4l(i).cell2(j).R = com_r;
        T4l(i).cell2(j).P = logP;
        f1 = EightCell_left_f(start(j):stop(j),7); %start and stop of each chromosome; column of cell 1
        f2 = EightCell_left_f(start(j):stop(j),8);
        r1 = EightCell_left_r(start(j):stop(j),7);
        r2 = EightCell_left_r(start(j):stop(j),8);
        [com_f, com_r, logP ] = Prob_Cal(f1, f2, r1, r2, b_4cell(4));
        T4l(i).cell3(j).F = com_f;
        T4l(i).cell3(j).R = com_r;
        T4l(i).cell3(j).P = logP;
        f1 = EightCell_left_f(start(j):stop(j),9); %start and stop of each chromosome; column of cell 1
        f2 = EightCell_left_f(start(j):stop(j),10);
        r1 = EightCell_left_r(start(j):stop(j),9);
        r2 = EightCell_left_r(start(j):stop(j),10);
        [com_f, com_r, logP ] = Prob_Cal( f1, f2, r1, r2, b_4cell(4));
        T4l(i).cell4(j).F = com_f;
        T4l(i).cell4(j).R = com_r;
        T4l(i).cell4(j).P = logP;
    end
end
% make to 2 cell stage
T2l = struct('cell1', struct, 'cell2', struct);
for i = 1:size(Premove,1)
    for j = 1:chr_num
        f1 = T4l(i).cell1(j).F;
        f2 = T4l(i).cell2(j).F;
        r1 = T4l(i).cell1(j).R;
        r2 = T4l(i).cell2(j).R;
        [com_f, com_r, logP ] = Prob_Cal( f1, f2, r1, r2, b_4cell(3));
        T2l(i).cell1(j).F = com_f;
        T2l(i).cell1(j).R = com_r;
        T2l(i).cell1(j).P = logP;
        f1 = T4l(i).cell3(j).F;
        f2 = T4l(i).cell4(j).F;
        r1 = T4l(i).cell3(j).R;
        r2 = T4l(i).cell4(j).R;
        [com_f, com_r, logP ] = Prob_Cal( f1, f2, r1, r2, b_4cell(3));
        T2l(i).cell2(j).F = com_f;
        T2l(i).cell2(j).R = com_r;
        T2l(i).cell2(j).P = logP;
    end
end
% make to 1 cell stage
T1l = struct;
for i = 1:size(Premove,1)
    for j = 1:chr_num
        f1 = T2l(i).cell1(j).F;
        f2 = T2l(i).cell2(j).F;
        r1 = T2l(i).cell1(j).R;
        r2 = T2l(i).cell2(j).R;
        [com_f, com_r, logP ] = Prob_Cal( f1, f2, r1, r2, b_4cell(2));
        T1l(i).cell1(j).F = com_f;
        T1l(i).cell1(j).R = com_r;
        T1l(i).cell1(j).P = logP;
    end
end
totPl = zeros(size(Premove,1),1);
% then for each tree sum all P
for i = 1:size(Premove,1) %save totPl for each case; then cross all the cases at once outside time for loop
    for j = 1:chr_num
        totPl(i,1) = totPl(i,1) + T4l(i).cell1(j).P + T4l(i).cell2(j).P + T4l(i).cell3(j).P + T4l(i).cell4(j).P ...
            +T2l(i).cell1(j).P + T2l(i).cell2(j).P + T1l(i).cell1(j).P;
    end
end
% right side 
Premove = PtotalRight;
T4r = struct('cell1', struct, 'cell2', struct,'cell3', struct,'cell4', struct);
for i = 1:size(Premove,1)
    EightCell_right_f(:,1:10) = SixteenCell_f(:,[1:2 PtotalRight(i,:)+2]);
    EightCell_right_r(:,1:10) = SixteenCell_r(:,[1:2 PtotalRight(i,:)+2]);
    for j = 1:chr_num
        f1 = EightCell_right_f(start(j):stop(j),3); %start and stop of each chromosome; column of cell 1
        f2 = EightCell_right_f(start(j):stop(j),4);
        r1 = EightCell_right_r(start(j):stop(j),3);
        r2 = EightCell_right_r(start(j):stop(j),4);
        [com_f, com_r, logP ] = Prob_Cal( f1, f2, r1, r2, b_4cell(4));
        T4r(i).cell1(j).F = com_f;
        T4r(i).cell1(j).R = com_r;
        T4r(i).cell1(j).P = logP;
        f1 = EightCell_right_f(start(j):stop(j),5); %start and stop of each chromosome; column of cell 1
        f2 = EightCell_right_f(start(j):stop(j),6);
        r1 = EightCell_right_r(start(j):stop(j),5);
        r2 = EightCell_right_r(start(j):stop(j),6);
        [com_f, com_r, logP ] = Prob_Cal( f1, f2, r1, r2, b_4cell(4));
        T4r(i).cell2(j).F = com_f;
        T4r(i).cell2(j).R = com_r;
        T4r(i).cell2(j).P = logP;
        f1 = EightCell_right_f(start(j):stop(j),7); %start and stop of each chromosome; column of cell 1
        f2 = EightCell_right_f(start(j):stop(j),8);
        r1 = EightCell_right_r(start(j):stop(j),7);
        r2 = EightCell_right_r(start(j):stop(j),8);
        [com_f, com_r, logP ] = Prob_Cal( f1, f2, r1, r2, b_4cell(4));
        T4r(i).cell3(j).F = com_f;
        T4r(i).cell3(j).R = com_r;
        T4r(i).cell3(j).P = logP;
        f1 = EightCell_right_f(start(j):stop(j),9); %start and stop of each chromosome; column of cell 1
        f2 = EightCell_right_f(start(j):stop(j),10);
        r1 = EightCell_right_r(start(j):stop(j),9);
        r2 = EightCell_right_r(start(j):stop(j),10);
        [com_f, com_r, logP ] = Prob_Cal( f1, f2, r1, r2, b_4cell(4));
        T4r(i).cell4(j).F = com_f;
        T4r(i).cell4(j).R = com_r;
        T4r(i).cell4(j).P = logP;
    end
end
% make to 2 cell stage
T2r = struct('cell1', struct, 'cell2', struct);
for i = 1:size(Premove,1)
    for j = 1:chr_num
        f1 = T4r(i).cell1(j).F;
        f2 = T4r(i).cell2(j).F;
        r1 = T4r(i).cell1(j).R;
        r2 = T4r(i).cell2(j).R;
        [com_f, com_r, logP ] = Prob_Cal( f1, f2, r1, r2, b_4cell(3));
        T2r(i).cell1(j).F = com_f;
        T2r(i).cell1(j).R = com_r;
        T2r(i).cell1(j).P = logP;
        f1 = T4r(i).cell3(j).F;
        f2 = T4r(i).cell4(j).F;
        r1 = T4r(i).cell3(j).R;
        r2 = T4r(i).cell4(j).R;
        [com_f, com_r, logP ] = Prob_Cal( f1, f2, r1, r2, b_4cell(3));
        T2r(i).cell2(j).F = com_f;
        T2r(i).cell2(j).R = com_r;
        T2r(i).cell2(j).P = logP;
    end
end
% make to 1 cell stage
T1r = struct; 
for i = 1:size(Premove,1)
    for j = 1:chr_num
        f1 = T2r(i).cell1(j).F;
        f2 = T2r(i).cell2(j).F;
        r1 = T2r(i).cell1(j).R;
        r2 = T2r(i).cell2(j).R;
        [com_f, com_r, logP ] = Prob_Cal( f1, f2, r1, r2, b_4cell(2));
        T1r(i).cell1(j).F = com_f;
        T1r(i).cell1(j).R = com_r;
        T1r(i).cell1(j).P = logP;
    end
end
% then for each tree sum all P
totPr = zeros(size(Premove,1),1);
for i = 1:size(Premove,1)
    for j = 1:chr_num
        totPr(i,1) = totPr(i,1) + T4r(i).cell1(j).P + T4r(i).cell2(j).P + T4r(i).cell3(j).P + T4r(i).cell4(j).P ...
            +T2r(i).cell1(j).P + T2r(i).cell2(j).P + T1r(i).cell1(j).P;
    end
end
totP = zeros(size(Ptotal,1),1); % the same rows as inputted Ptotal 
for i = 1:size(Ptotal,1) %going case by case
    left_side = Ptotal(i,1:8);
    right_side = Ptotal(i,9:16);
    % find the row in Ptotalleft or right
    index_left = ismember(PtotalLeft,left_side,'rows'); % should never have more than 1 row 
    index_right = ismember(PtotalRight,right_side,'rows');
    totP(i,1) = totPl(index_left) + totPr(index_right);
end
totP2 = -totP;
totP2 = totP2 - max(totP2) + 100; % normalized max to 100 such that the top is only 100 so that exp would not explode 
% this is in the ln scale
expP = exp(totP2);
expP_sum = sum(expP);
expP_prob = expP /expP_sum;
expP_prob(1:size(Ptotal,1),2) = 1:size(Ptotal,1);
expP_sort = sortrows(expP_prob,'descend');
expP_sort_top = expP_sort(1:max_num,:);
end
```
