## Supplementary material for "A probabilistic framework for cellular lineage reconstruction using single-cell 5-hydroxymethylcytosine sequencing": Code (scPECLR): sc_PECLR_test.m

```
%% This file is the test file for scTRACE
% choose the path of all the workspaces and functions
addpath('/Users/admin/Documents/MATLAB/Algorithm/scPECLR/');

%% 8-cell data
load('test8cell_embryo.mat');
[fin_result_8cell, tree_result_8cell, Ptotal_8cell] = scPECLR(EightCell_f,EightCell_r);

%% 16-cell data 
load('test16cell_embryo.mat');
[fin_result_16cell, tree_result_16cell, Ptotal_16cell] = scPECLR(SixteenCell_f,SixteenCell_r);
```
