## Supplementary material for "A probabilistic framework for cellular lineage reconstruction using single-cell 5-hydroxymethylcytosine sequencing": Code (scPECLR): scPECLR.m

```
function [fin_result, tree_result, Ptotal] = scTRACE(forward_5hmC,reverse_5hmC)
% input both plus (forward) and minus (reverse) strands (in 1/0 format) with no overlap. The first two
% columns are the chromosomes and the bins, the rest of the columns are
% 5hmC within each cell (8 or 16 cells). Both input must have the same size.
% fin_result - row = iteration number, 1st column - tree predicted, 2/3/4 values of b used in the iteration
% tree_reult - row = iteration number, 1st column - tree predicted, 2nd its
% associated prob, 3/4 and 5/6 are the second and third most likely trees
% respectively.
% Ptotal - tree topologies considered 
% 8 cells give top 3 trees, 16 cells give top 10 trees (can change but 8cells should not have more than 9 trees) 
Cell_f = forward_5hmC;
Cell_r = reverse_5hmC;
min_b = 0.01; %set minimum b for iterative calculation
iter_max = 10;
cell_num = size(Cell_f,2)-2;
if cell_num == 8
    tree_max = 3; 
    [Ptotal, ~, ~, totbin] = OSS_8(Cell_f ,Cell_r);
    [tree_result,fin_result] = Prediction_8(Cell_f ,Cell_r, Ptotal, totbin, min_b, iter_max, tree_max);
elseif cell_num == 16
    tree_max = 10;
    [Ptotal, ~, ~, totbin] = OSS_16(Cell_f, Cell_r);
    [tree_result,fin_result] = Prediction_16(Cell_f ,Cell_r, Ptotal, totbin, min_b, iter_max, tree_max);
else
    error('cell number is not 8 or 16');
end
end
```
